# Engineered Nanobodies Bind Theranostic Main Group Metals

**DOI:** 10.1101/2024.09.26.615111

**Authors:** Pritha Ghosh, Lani J. Davies, Christoph Nitsche

## Abstract

Targeted theranostics heavily rely on metal isotopes conjugated to antibodies. Single-domain antibodies, known as nanobodies, are much smaller in size without compromising specificity and affinity. The conventional way of conjugating metals to nanobodies involves non-specific modification of amino acid residues with bifunctional chelating agents. We demonstrate that mutagenesis of a single residue in a nanobody creates a triple cysteine motif that selectively binds bismuth which is, for example, used in targeted alpha therapy. Two mutations create a quadruple cysteine mutant specific for gallium and indium used in positron emission tomography and single-photon emission computed tomography, respectively. Labelling is quantitative within a few minutes. The metal nanobodies maintain structural integrity and stability over weeks, resist competition from endogenous metal binders like glutathione, and retain functionality.

---

The integration of monoclonal antibodies with metals is crucial for theranostic applications (*1-3*). While the antibody typically targets a specific biomolecule, such as a cancer cell receptor, the metal serves as a probe for imaging purposes or selectively destroys target cells through radiation. For example, the monoclonal antibody trastuzumab, which binds to the human epidermal growth factor receptor 2 (HER2), can be labeled with the radioactive isotopes ^68^Ga or ^111^In for imaging purposes using positron emission tomography (PET) or single-photon emission computed tomography (SPECT), respectively (*4, 5*). Combination of trastuzumab with the radioactive isotope ^213^Bi allows the selective destruction of HER2-positive cancer cells using targeted alpha therapy (TAT) (*6*). Further applications can be found in chemical biology, where antibodies tagged with isotopes such as ^209^Bi are used for single-cell mass cytometry (*7*).

While antibodies continue to dominate sales and clinical approvals in biopharmaceuticals (*8*), their large size (∼150 kDa) limits tumor tissue penetration, and their glycosylation necessitates production in mammalian cells. Single-domain antibodies (sdAb), better known as nanobodies, are an emerging class of antigen-binding fragments originating from camelids and cartilaginous fishes (*9*). While they exhibit affinity and selectivity similar to conventional antibodies, their much smaller size (12–15 kDa) allows for better tissue penetration and they can be recombinantly expressed from bacterial culture (*10*). Attributed to their refolding capability, nanobodies possess extraordinary stability, including a prolonged shelf life at 4 °C, resistance to proteolysis, and tolerance to high temperatures of 60–80 °C, a broad pH range of 3 to 9, and denaturants (*11*). These properties position nanobodies as promising candidates for theranostics and applications in chemical biology (*12*). It is hence no surprise that, like trastuzumab, nanobodies can equally target HER2 and have been combined with ^213^Bi in a TAT strategy (*13*) and with ^68^Ga or ^111^In for PET and SPECT imaging, respectively (*14-16*).

The conventional strategy for conjugating antibodies or nanobodies to metals uses bifunctional linkers (*17, 18*). One part of these linkers reacts with lysine or cysteine residues in the protein, while the other part contains a metal chelator (Fig. 1A). Though specific modification of cysteine residues is possible through mutagenesis (*16*), site-selective modification of lysine residues remains challenging (*19*). Typically, residues are randomly modified, resulting in heterogeneous conjugates and partial inactivation by altering the antigen-binding site. In addition to challenges with harsh, incomplete and lengthy tagging processes (*17, 20*), conventional chelators are large and highly charged, impacting on the properties of the conjugate. This is particularly concerning for nanobodies, where the bifunctional linker significantly changes the overall mass and charge. It would therefore be attractive to attach the metal directly to the nanobody and hence abolish bifunctional linkers, which is pioneered in this study (Fig. 1B).

**Fig. 1.**
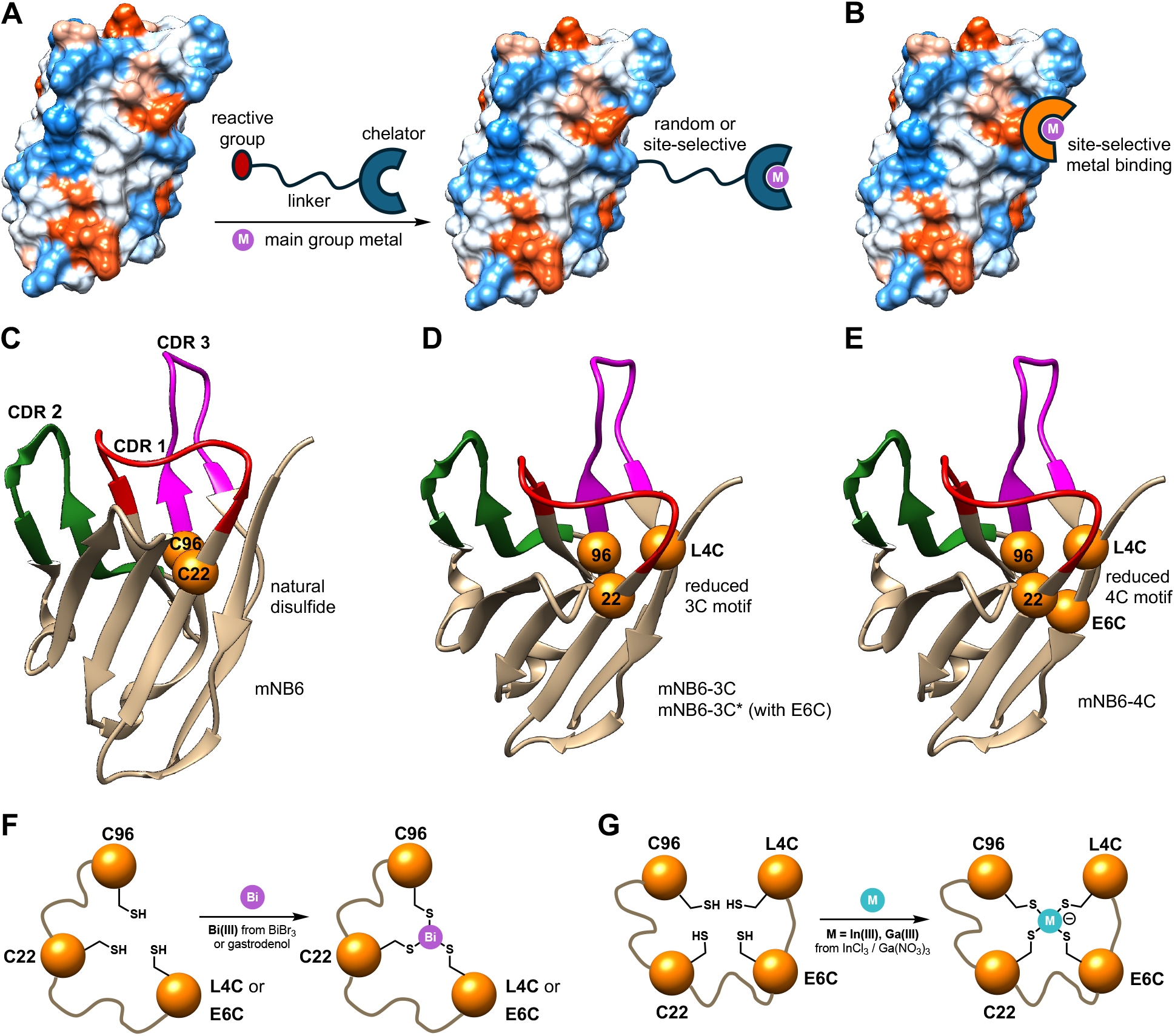
Nanobody design and metal binding. (**A**) Conventional strategy to label nanobodies with bifunctional linkers. (**B**) This study introduces a metal-binding site directly inside the nanobody. (**C**) Structure of a wildtype nanobody (anti-SARS-CoV-2 spike nanobody mNb6; pdb: 7KKJ). The conserved disulfide bond (C22, C96) and the three complementarity-determining regions (CDRs) are indicated. (**D**) A single mutation, L4C, generates a triple cysteine motif. (**E**) An additional mutation, E6C, generates a quadruple cysteine motif with tetrahedral geometry. A single E6C mutations generates an alternative triple cysteine motif. (**F**) Bismuth(III) binding to the primed triple cysteine motif. (**G**) Indium(III) and gallium(III) binding to the primed quadruple cysteine motif.

## Designing binding motifs for main group metals in nanobodies

As the three main group elements gallium, indium and bismuth cover the major applications in imaging and radiotherapy with PET, SPECT and TAT, we set out to generate a common binding site using cysteine residues based on their thiophilic properties (*21*). We founded our design on the disulfide bond between C22 and C96 in nanobodies (Fig. 1C), which, while highly conserved, is neither essential for stability and activity (*22*) nor close to binding epitopes (*23-25*). Our previous work on peptides demonstrated that three cysteine residues form stable complexes with Bi(III) (*26*), hence we introduced a third cysteine residue within 4–8 Å of the native disulfide in nanobodies to create a triple cysteine motif for bismuth binding (Fig. 1, D and F). Because In(III) and Ga(III) form negatively charged indates and gallates with cysteine-based ligands (*27*), we also designed a nanobody mutant with two additional cysteine residues adjacent to the native disulfide, creating a quadruple cysteine motif for gallium and indium recognition (Fig. 1, E and G).

## Optimizing disulfide reduction and metal uptake

As an initial model system, we chose a previously reported nanobody, mNb6, raised against the receptor binding domain (RBD) of the spike protein of SARS-CoV-2 (*24*). We conducted periplasmic expression in *E. coli* of the wildtype (*24*), the L4C (mNb6-3C) and E6C (mNb6-3C*) single mutants, and the L4C/E6C (mNb6-4C) double mutant (Fig. 1, C to E, Fig. 2H). As expected, the wildtype mNb6 and mutants mNb6-3C and mNb6-3C* contain one disulfide bond after expression, while mNb6-4C displayed two disulfides (Fig. 2, A, B and C). To coordinate Bi(III) to mNb6-3C, we first attempted to reduce the disulfide bond in large excess of the reducing agent tris(2-carboxyethyl)phosphine (TCEP) at room temperature. Neither co-incubation with the water-soluble Bi(III) drug gastrodenol nor its addition at a later point resulted in any significant formation of the mNb6-3C-Bi conjugate (Fig. 2I). As mass spectrometry (MS) confirmed the retention of the disulfide bond after TCEP treatment (fig. S1), we concluded that the disulfide is inaccessible, and that partial unfolding would be required. Chemical denaturation induced by guanidine resulted in detectable, though incomplete modification (fig. S2). Informed by the denaturation temperature (Fig. 2F), short heating to 50 °C quantitatively yielded the bismuth-modified nanobodies mNb6-3C-Bi and mNb6-3C*-Bi, as confirmed by native MS (Fig. 2, A, B and I).

**Fig. 2.**
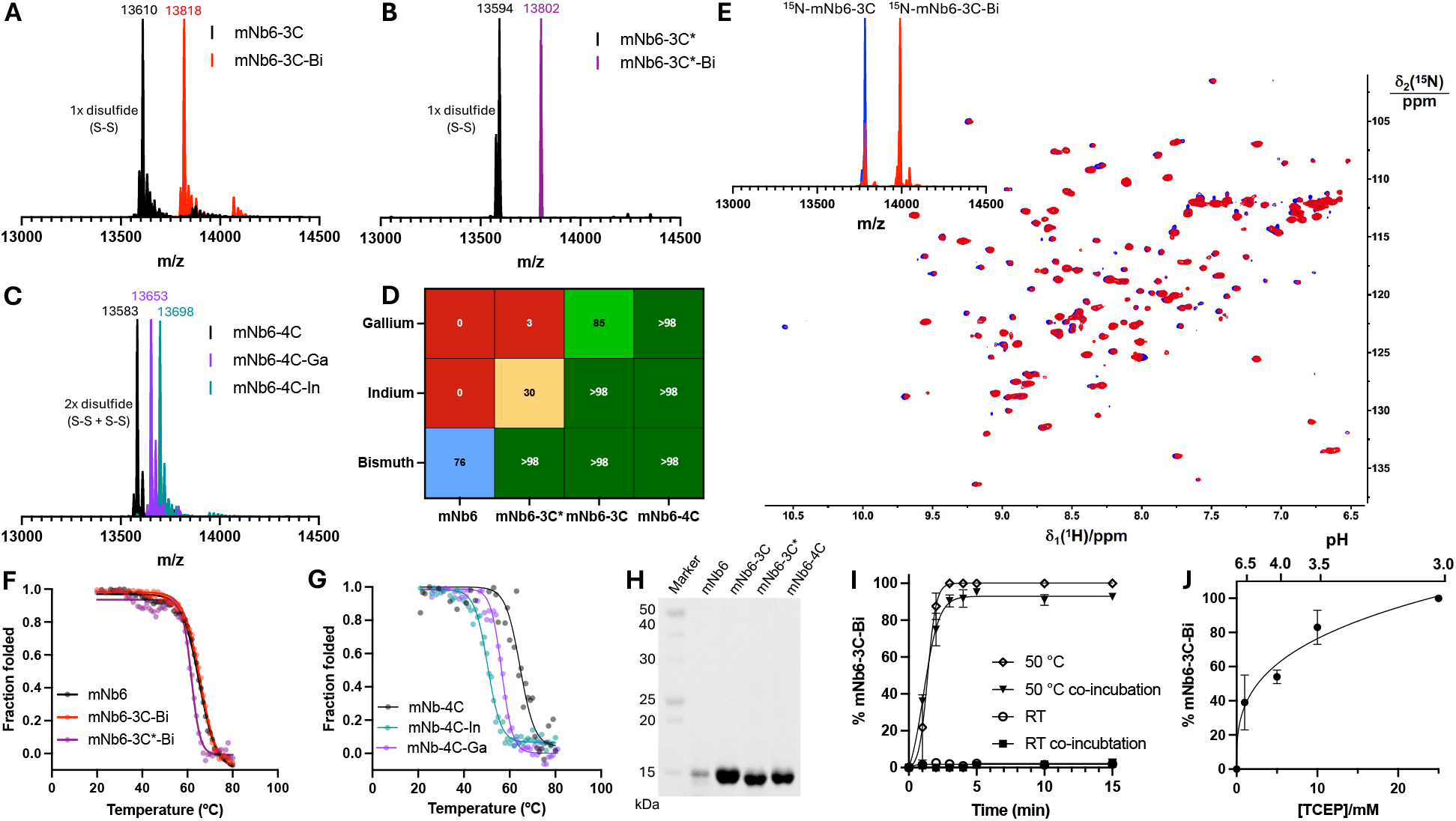
Disulfide reduction and metal modification of nanobodies. (**A**–**C**) Native mass spectrometry (MS) performed in 100 mM ammonium acetate pH 7.0 for mNb6-3C, mNb6-3C* and mNb6-4C before and after modification with Bi(III), Ga(III) or In(III). Nanobodies were reduced with 25 mM TCEP for 15 min at 50 °C (mNb6-3C and mNb6-3C*) or 60 °C (mNb6-4C) prior addition of the metal reagent. (**D**) Uptake (%) of Ga(III), In(III) and Bi(III) by nanobodies containing two, three or four cysteine residues after reduction with 25 mM TCEP for 15 min at 55 °C (mNb6), 50 °C (mNb6-3C and mNb6-3C*) and 60 °C (mNb6-4C) determined by native MS. (**E**) Superimposition of 800 MHz [^15^N,^1^H]-HSQC NMR spectra of 150 μM solutions of mNb6-3C (blue) and mNb6-3C-Bi (red) in 20 mM MES pH 6.5, 150 mM NaCl, 10% D_2_O. The mNb6-3C-Bi sample additionally contains 10 mM TCEP. Native MS spectra of both samples are superimposed. (**F**,**G**) Thermal denaturation curves determined by circular dichroism (CD) spectroscopy in 20 mM phosphate buffer pH 7.4 for mNb6 (T_m_ = 65 °C), mNb6-3C-Bi (T_m_ = 65 °C), mNb6-3C*-Bi (T_m_ = 62 °C), mNb6-4C (T_m_ = 65 °C), mNb6-4C-Ga (T_m_ = 56 °C), and mNb6-4C-In (T_m_ = 51 °C). (**H**) SDS-PAGE of mNb6 nanobody constructs after periplasmic expression from *E. coli* and His-tag affinity purification. (**I**) Time-dependent reduction and modification of mNb6-3C with 25 mM TCEP and 5 equiv. of Bi(III) from gastrodenol (bismuth tripotassium dicitrate) at room temperature (RT) or 50 °C. Bi(III) was either co-incubated or added after the reduction period as indicated, and uptake was monitored by native MS. (**J**) Bi(III) uptake of mNb6-3C monitored by native MS after incubation for 15 min at 50 °C with 5 equiv. of gastrodenol depending on TCEP concentration and pH.

While the 50 °C heat shock proved crucial for quantitative disulfide reduction, it resulted in the loss of up to 75% of soluble nanobody after 15 minutes at pH 7.5 (*28*). Contrary, at pH 3 most of the nanobody remained soluble at this temperature, highlighting the extraordinary stability at pH extremes (table S5, fig. S3). Simultaneous increase of the TCEP concentration and decrease of pH resulted in fastest disulfide reduction and absence of protein precipitation during heat shock (Fig. 2J). While treatment with BiBr_3_ is possible (fig. S4 and S5) (*26*), we observed quantitative modification within only 3 minutes using water-soluble gastrodenol (Fig. 2I) (*29*). The thermal denaturation curves of mNb6-3C-Bi and mNb6-3C*-Bi were found to be almost identical to the wildtype mNb6 (Fig. 2F), indicating that the bismuth-bridged cysteines mimic the native disulfide bond without impacting overall stability. Despite the likelihood of partial unfolding during the heat and pH shocks, circular dichroism (CD) spectroscopy indicated correctly folded nanobody post treatment (fig. S6). To further confirm that the tertiary structure remains intact throughout modification, we conducted nuclear magnetic resonance (NMR) spectroscopy with uniformly ^15^N-labelled mNb6-3C (Fig. 2E, fig. S7 and S8). Direct reduction in the NMR buffer at pH 6.5 for 15 minutes at 50 °C resulted in only 73% transformation into ^15^N-mNb6-3C-Bi (Fig. 2E), confirming that fast and quantitative modification is best achieved at pH 3. Superimposition of NMR spectra reveal only minimal peak perturbations upon bismuth binding (Fig. 2E). Bound and unbound species are in slow exchange on the NMR time scale, indicating a kinetically stable complex.

Applying our heat-and-pH-shock procedure to the quadruple cysteine motif in mNb6-4C, we were able to completely modify the nanobody with Ga(III) and In(III), using water-soluble Ga(NO_3_)_3_ and InCl_3_, respectively (Fig. 2C). It is unsurprising that full complexation occurred with indium due to its high thiophilicity (*21*). Gallium, however, as a hard Lewis acid is more drawn toward oxygen and nitrogen ligands; hence, full complexation of Ga(III) by mNb6-4C surprised us. High-resolution native MS confirmed that all four cysteine thiols are deprotonated, engaging in the formation of negatively charged indate [InSR_4_]^−^ and gallate [GaSR_4_]^−^ complexes (Fig. 1G, Fig. 3, F and G). We quantified and comprehensively compared uptake of Bi(III), In(III) and Ga(III) by nanobodies comprising two (mNb6), three (mNb6-3C and mNb6-3C*) and four (mNb6-4C) cysteine residues after a 15-minute heat-and-pH-shock using native MS (Fig. 2D, fig. S9 – S15). Only bismuth binds to the wildtype 2C motif, although not quantitatively unless fully denatured (fig. S10). The 3C motif quantitatively binds bismuth and indium, while gallium can only be fully complexed by the 4C motif.

**Fig. 3.**
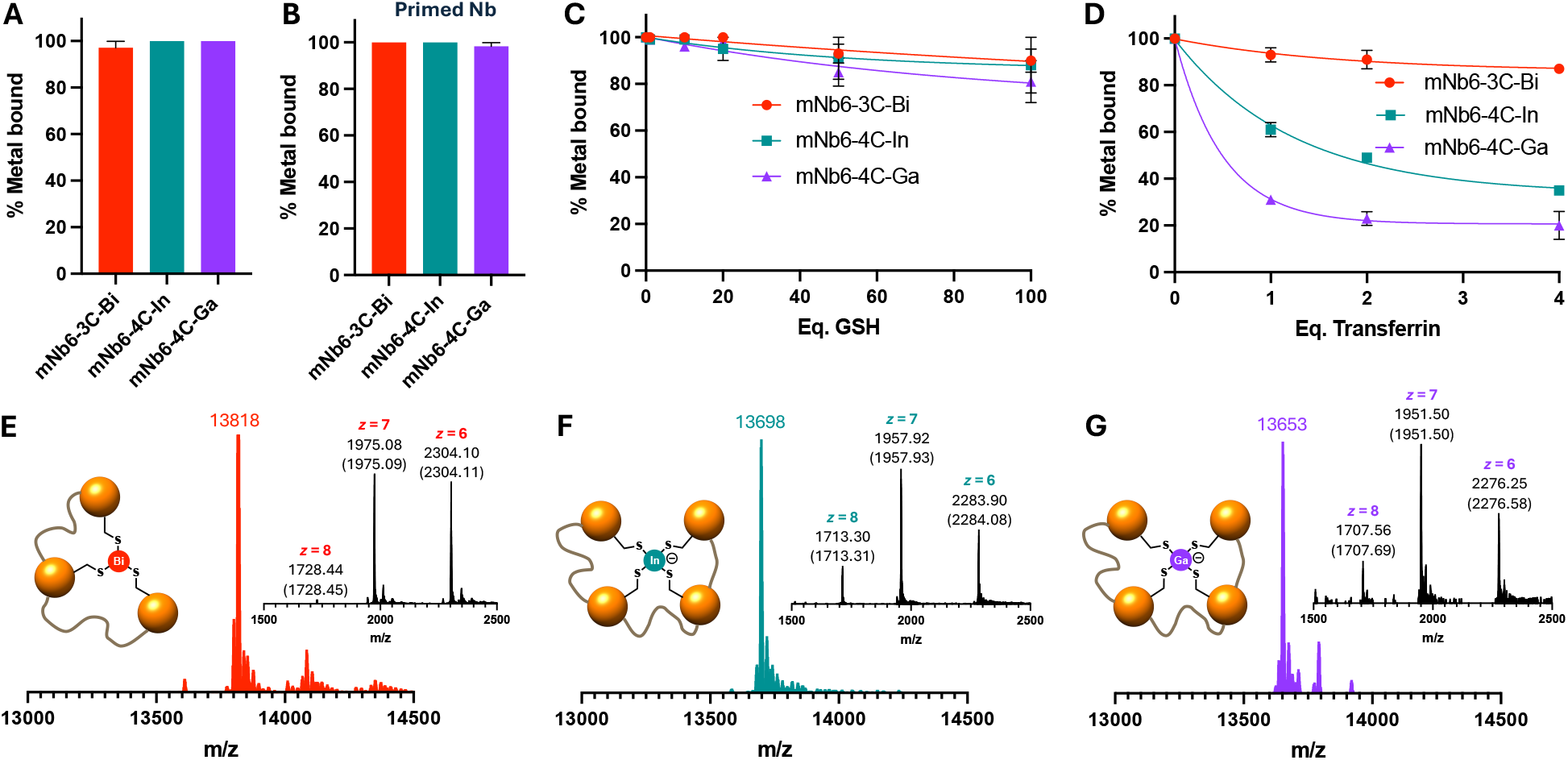
Main group metal nanobody stability. (**A**) Retention of Bi(III), In(III) and Ga(III) in engineered mNb6 variants after two weeks of storage at 4 °C in 100 mM ammonium acetate, pH 7.0. (**B**) Uptake of Bi(III), In(III) and Ga(III) by engineered mNb6 variants two weeks after reduction and storage at 4 °C in 20 mM Tris-HCl, pH 7.5, 150 mM NaCl, 25 mM TCEP. (**C**,**D**) Retention of Bi(III), In(III) and Ga(III) bound to engineered mNb6 variants [50 μM for (C), 25 μM for (D)] in presence of glutathione (C) and apo-transferrin (D) after 1 h incubation at 25 °C in 100 mM ammonium acetate, pH 7.0. (**E**–**F**) Original and deconvoluted high-resolution native MS spectra of mNb6-3C-Bi, mNb6-4C-In and mNb6-4C-Ga after 18 h dialysis at 4 °C in 20 mM HEPES, pH 7.8, 150 mM NaCl. Calculated m/z ratios for the indicated species are given in brackets.

## Stability of main group metal nanobodies

After identifying suitable binding motifs for bismuth, indium and gallium in the model nanobody mNb6, we set out to study the stability of these metal-nanobody conjugates (Fig. 3). Following modification and buffer exchange, all three major conjugates, mNb6-3C-Bi, mNb6-4C-In and mNb6-4C-Ga, remain fully intact if stored for two weeks at 4 °C (Fig. 3A, fig. S16 – S23). To provoke metal dissociation and fully eliminate the possibility that slight metal excess or reagents like TCEP could impact our stability assessment, we dialyzed (1000:1 *v*/*v*) the nanobody-metal conjugates for 18 h at 4 °C and observed that after this period >98% of the metal remained bound (Fig. 3, E, F and G). We further co-dialyzed mNb6-4C-In and mNb6-4C-Ga, which share the identical construct, and only identified <1% cross-contamination of indium and gallium after 18 h (Fig. 3, F and G) in each sample, demonstrating extremely slow metal dissociation sufficient for clinical applications.

As it can be beneficial for clinical set-ups dealing with radioactive metals with limited half-lives, such as ^213^Bi (*T*_1/2_ = 46 min) used in TAT (*30*), to add the metal at a very late stage, we explored if reduction and metal uptake can be divided into two distinct steps, separated by days if not weeks. After standard reduction procedure, we kept the ‘primed’ nanobodies at 4 °C and added the metals after different time points (fig. S24, S25). Even after two weeks of storage at 4 °C, we observed >98% metal uptake across all nanobodies and metals tested (Fig. 3B), highlighting that primed nanobodies can be transported to the site where both the reactor and patient are present.

## Competition with endogenous metal binders

In order to further assess the complex stability of mNb6-3C-Bi, mNb6-3C*-Bi, mNb6-4C-In and mNb6-4C-Ga, we performed competition experiments with glutathione (Fig. 3C, fig. S26) and human transferrin (Fig. 3D, fig. S26). Glutathione (GSH) is the most abundant thiol in human cells (1–2 mM), playing a key role in maintaining the reducing environment, detoxification, and metal homeostasis (*31*). Though the primary function of transferrin is to bind and transport iron, it also binds various heavy metals to support detoxification (*32*). While the thiol in GSH can be considered a soft ligand, transferrin is rather a hard ligand with nitrogen and oxygen donors in its two binding sites comprised of tyrosines, histidine, arginine and aspartate.

The three nanobody-metal conjugates proved remarkably stable against GSH after 1 h of incubation (Fig. 3C). At 5 mM GSH, which even exceeds most cellular levels, 80–90% of the nanobody molecules retained the metal, as determined by native MS. At a more realistic concentration of 50 μM, which still exceeds typical plasma levels (*33*), >98% of nanobody-metal conjugates remained intact (Fig. 3C). The complex stability constant log*K* between Bi(III) and three GSH ligands, [Bi(GS)_3_], has previously been determined by displacement of EDTA to be 30.1 at pH 7.3 (*34*). The fact that even 100 equivalents of GSH displace less than 10% Bi(III) from mNb6-3C-Bi (Fig. 3C) demonstrates that the thermodynamic complex stability is within the range of conventional as well as contemporary Bi(III) complexing agents (*35, 36*).

The transferrin competition experiments yielded a more distinct picture with pronounced differences between bismuth, indium and gallium (Fig. 3D). All three elements are known to bind to human transferrin with similar stability constants (*37-39*), and a crystal structure for the bismuth complex has been reported (*40*). To avoid any unwanted interference from iron, we opted to use apo-transferrin, which has both metal-binding sites unoccupied, unlike transferrin in healthy humans, which is typically 30% saturated (*32*). After 1 h incubation with 50 μM apo-transferrin, mNb6-3C-Bi maintains >90% bismuth saturation. This concentration exceeds typical plasma levels of transferrin and relates to a 4:1 ratio of available metal-binding sites in transferrin and the nanobody. While the impact on bismuth binding was marginal, transferrin had a much more pronounced effect on indium and gallium bound to mNb6-4C (Fig. 3D). Whilst at 50 μM apo-transferrin, mNb6-4C still showed 50% saturation with indium, it only displayed 25% gallium saturation. These observations align with the hard and soft acids and bases (HSAB) theory. Gallium is a harder Lewis acid than indium and the engineered nanobody is a softer ligand than transferrin. Overall, our GSH and transferrin competition experiments indicate the following ranking with respect to inertness of the nanobody-metal conjugates: Bi >> In > Ga.

## Metal-nanobody conjugates remain fully functional

We further assessed whether the engineered nanobodies still bind their target proteins after modification with the investigated main group metals. After confirming full modification by native MS, we immobilized mNb6-3C-Bi, mNb6-4C-In and mNb6-4C-Ga on a CM5 chip and conducted surface plasmon resonance (SPR) experiments by flushing the RBD of the SARS-CoV-2 spike protein over the chip surface (fig. S27). These experiments confirmed fully functional nanobodies with dissociation constants (*K*_D_) between 17 and 34 nM (Fig. 4, A, B and C), demonstrating tight binding of the bismuth, indium and gallium nanobodies to their target (fig. S28).

**Fig. 4.**
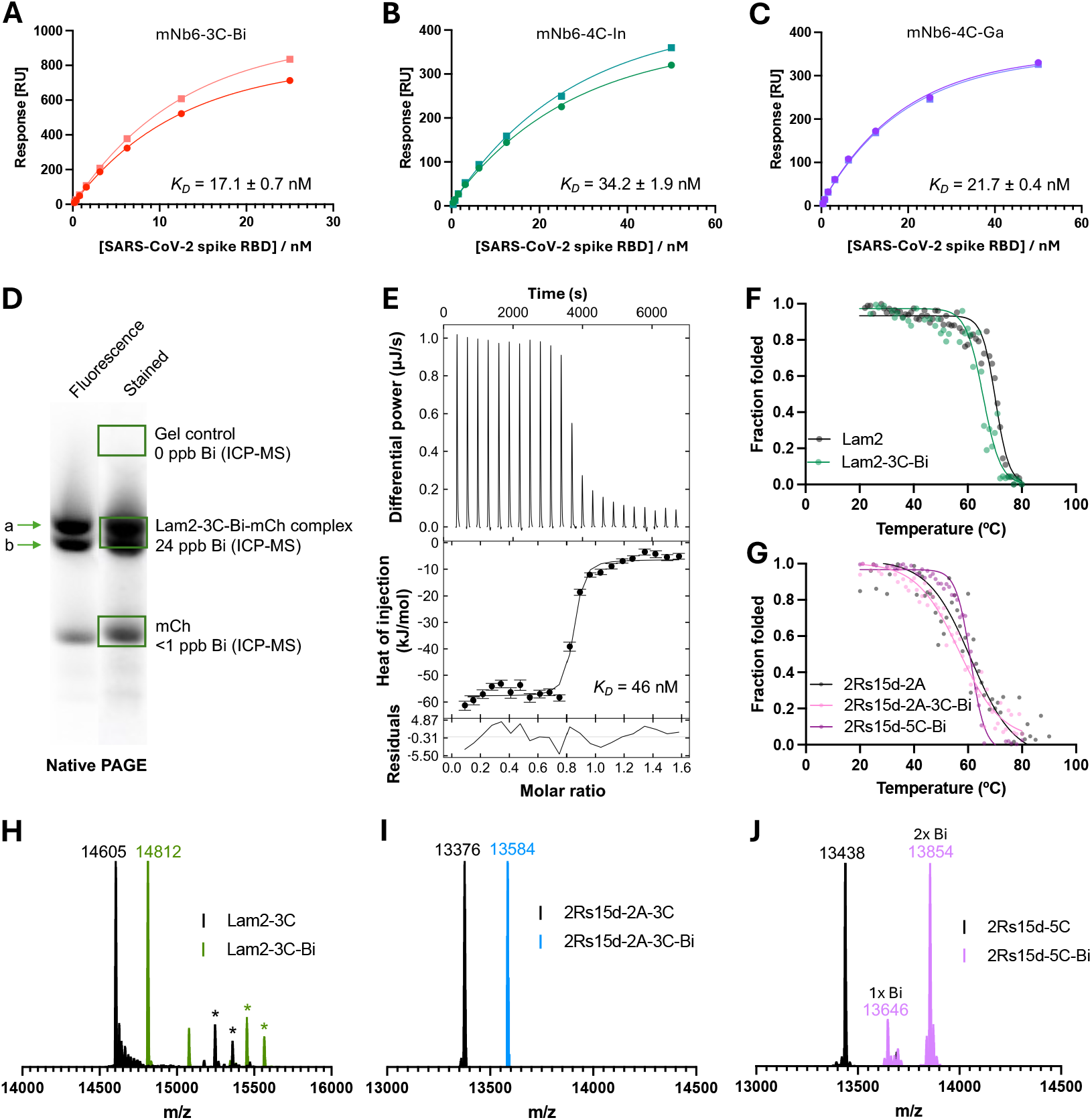
Functionality and scope of metal nanobodies. (**A**–**C**) Binding curves and dissociation constants of the receptor binding domain (RBD) of SARS-CoV-2 and mNb6-3C-Bi (A), mNb6-4C-In (B) and mNb6-4C-Ga (C) determined by surface plasmon resonance (SPR). (**D**) Native PAGE of Lam2-Bi nanobody and mCherry (mCh). Excised gel parts are indicated by green boxes and their bismuth content, determined by ICP-MS, is indicated as parts per billion (ppb). Two bands for the complex are a result of mNb6-3C being excreted with (a) and without (b) part of its N-terminal leader sequence. (**E**) Isothermal titration calorimetry (ITC) of Lam2-Bi and mCh with indicated dissociation constant (*K*_D_). (**F**,**G**) Thermal denaturation curves determined by CD spectroscopy in 20 mM phosphate buffer pH 7.4 for Lam2 (T_m_ = 70 °C), Lam2-3C-Bi (T_m_ = 66 °C), 2Rs15d-2A (T_m_ = 61 °C), 2Rs15d-2A-3C-Bi (T_m_ = 58 °C), and 2Rs15d-5C-Bi (T_m_ = 61 °C). (**H**) Superimposed native MS spectra of Lam2-3C and Lam2-3C-Bi. Species of higher m/z ratio that correspond to a partially cleaved N-terminal leader sequence are indicated with an asterisk (*). (**I**,**J**) Superimposed intact MS spectra of 2Rs15d-2A-3C (I) and 2Rs15d-5C (J) in presence and absence of Bi(III).

In order to further expand the scope and validate that the approach is broadly applicable to nanobodies raised against various targets, we selected the nanobody Lam2 binding to the red fluorescent protein mCherry (mCh) frequently used in chemical biology (*25, 41*). Following from the work with mNb6-3C, we introduced the same L4C mutation in Lam2-3C. Periplasmic expression resulted in species where the N-terminal leader sequence was either fully or only partially cleaved (fig. S29). Informed by the thermal denaturation curve (Fig. 4F), the heat-and-pH-shock method led to correctly folded (fig. S30) Lam2-3C-Bi with >98% bismuth saturation as confirmed by native MS (Fig. 4H, fig. S31 and S32). Isothermal titration calorimetry (ITC) of Lam2-3C-Bi with mCherry confirmed fully functional modified nanobody with a *K*_D_ of 46 nM exceeding the wildtype Lam2 (Fig. 4E, table S6, fig. S33). To demonstrate beyond any doubt that bismuth remains conjugated when the nanobody binds to its target protein, we conducted native PAGE with Lam2-3C-Bi and mCherry, followed by inductively coupled plasma MS (ICP-MS) analysis of individual gel bands (Fig. 4D). This combination of techniques confirmed not only the formation of the mCherry-Lam2-3C-Bi complex under native conditions, but also proved the presence of bismuth only in the complex and not in mCherry itself (Fig. 4D, table S7).

Finally, we evaluated another therapeutically relevant nanobody, 2Rs15d, that binds to the HER2 receptor overexpressed in some cancer cells (*23, 42*). This nanobody contains an additional non-canonical disulfide bond between CDR 1 and CDR 3 (C32 and C98) that is typical for single-domain antibodies of camelid origin. While without impact on functionality, this optional disulfide bond can contribute to overall nanobody stability (*43*). Hence, we first investigated an L4C mutant of 2Rs15d with both disulfide bonds retained, resulting in a total of five cysteine residues in 2Rs15d-5C for which we confirmed correct folding (fig. S34) and thermal denaturation above 60 °C (Fig. 4G) by CD spectroscopy. The usual heat-and-pH-shock and treatment with 5 equivalents of gastrodenol yielded 2Rs15d-5C-Bi with two bound bismuth atoms as the dominant species (Fig. 4J). The fact that the extra disulfide bond is capable of binding Bi(III) aligns with our mNb6 experiments, where we observed 76% bismuth saturation for soluble protein (Fig. 2D).

While 2Rs15d-5C demonstrates the opportunity for increasing the overall bismuth load per nanobody, it must be noted that the 2C-bismuth conjugate appears to be much more labile than our engineered 3C bismuth trap, as indicated by the presence of the minor single-bismuth species in 2Rs15d-5C-Bi (Fig. 4J). Therefore, we additionally explored a 2Rs15d-2A-3C mutant in which the non-canonical disulfide bond was mutated to two alanine residues. We also expressed the wildtype 2Rs15d-2A lacking the extra cysteine residue and confirmed folding and denaturation parameters in comparison to 2Rs15d-2A-3C (Fig. 4G, fig. S34 and S35). Like mNb6-3C and Lam-2-3C, 2Rs15d-3C could be fully saturated with one bismuth atom using our proven heat-and-pH-shock procedure (Fig. 4I). Overall, these experiments showcase that the L4C mutation reliably generates a common motif for main group metal binding across a variety of nanobodies.

## Conclusion

We have shifted the paradigm on how single-domain antibodies and main group metals are ligated for radiopharmaceutical applications and beyond. Metal-binding motifs can be constructed with nature’s toolbox of canonical amino acids, eliminating the need for expensive bifunctional chemical linkers. A single residue mutation is sufficient to create a site that binds Bi(III) as strongly as conventional chelators, withstanding competition by major cellular and plasma chelators like glutathione and transferrin. Aligning with the expected chemistry, our data demonstrate that the 3C motif is optimal for bismuth binding, while the 4C motif appears optimal for indium and gallium. In contrast to bifunctional chelators that expose the metal and separate it from the targeting biomolecule, our approach buries the metal inside the hydrophobic core of the nanobody, preventing it from dissociating. The approach is fully compatible with biotechnological production of nanobodies, as chemical modifications are no longer necessary. Separation of reduction and metal uptake enables production and application at different sites. We therefore anticipate broad applications across targeted radiotherapy, diagnostics and chemical biology.

## Acknowledgments

We gratefully acknowledge the Australian Research Council for funding support, including a Discovery Project (DP230100079) and a Future Fellowship (FT220100010). We thank Dr. Doug Lawes (NMR Suite, Research School of Chemistry, ANU) for his invaluable assistance with NMR acquisition. We also extend our appreciation to the staff of the Joint Mass Spectrometry Facility (ANU) for their guidance in training, sample preparation, and instrument handling. We thank Dr. Shouvik Aditya (Research School of Biology, ANU) for his support in SPR training and analysis and Dr. Yang Wu (Research School of Earth Sciences, ANU) for his support in ICP-MS sample preparation and instrumentation.

