## Supplementary Material for "Engineered Nanobodies Bind Theranostic Main Group Metals"

**The PDF file includes:**

Materials and Methods
Figs. S1 to S35
Tables S1 to S7
References

**Materials and Methods**

Materials

All plasmids encoding an N-terminal periplasmic leader sequence and C-terminal $His_6$ tag were obtained from Twist Bioscience, USA. The $His_6$ tagged receptor binding domain (aa 319–541) of the SARS-CoV-2 spike protein (RP-87678) was purchased from Invitrogen, Thermo Fisher Scientific, USA. Recombinant human Her2/ERBB2 protein (ECD, $His_6$ tag, 10004-H08H) was purchased from Sino Biological, China. Gastrodenol (bismuth tripotassium dicitrate), glutathione (~98% reduced), Tris, and IPTG were purchased from AK Scientific, USA. Indium(III) chloride hexahydrate, gallium(III) nitrate hydrate, ammonium acetate (LC-MS grade), imidazole, and human apo-transferrin were purchased from Sigma Aldrich, Australia. TCEP hydrochloride was purchased from AmBeed, USA. HEPES (free acid) and MES (free acid) monohydrate were purchased from Astral Scientific, Australia. $^{15}N$-labelled ammonium chloride (>99%) was purchased from Martek Isotopes LLC, USA. Kanamycin was purchased from AG Scientific, USA. Other chemicals and buffer compositions, such as sodium, potassium, and magnesium salts as well as EDTA were purchased from Sigma Aldrich, Australia. LC-MS grade solvents were purchased from Fisher Scientific, Australia. Bacterial media components were purchased from Gibco, Thermo Fisher Scientific, USA. Series S CM5 chip, amine coupling kit, and HisTrap™ (5 mL) columns were purchased from Cytiva, USA. Pre-cast SDS-PAGE and native-PAGE gels (Novex Bis-Tris Plus Mini Protein Gels) were purchased from Invitrogen, Thermo Fisher Scientific, USA. Unstained protein standard, broad range (10-200 KDa) was purchased from New England Biolabs, USA.

Expression and purification of nanobodies

Nanobody (Nb) sequences with N-terminal periplasmic leader sequence (MKYLLPTAAAGLLLLAAQPAMA) and C-terminal $His_6$ tag were cloned into a pET-29b(+) expression vector (Twist Bioscience, USA) and transformed into *E. coli*. BL21 (DE3). The expression protocol was followed as described by Schoof et al. (*24*) with minor changes. Isolated single colonies were grown in Luria Broth (LB) overnight, transferred to Terrific Broth and grown at 37 °C with gentle shaking until the OD reached 0.6–0.8. This was followed by induction of overexpression using 1 mM IPTG for 18–20 h at 25 °C. The *E. coli* cells were harvested (centrifuged at 5000 × *g*, 30 minutes, 4 °C) and resuspended in lysis buffer (200 mM Tris, pH 8.0, 500 mM sucrose, 0.5 mM EDTA) for 30 minutes on ice. This was followed by a 45-minute osmotic shock with a two-fold volume addition of cold water on ice with occasional mixing. 150 mM NaCl, 2 mM $MgCl_2$ and 40 mM imidazole were added to the lysate before centrifugation at 17,000 × *g* for 30 minutes at 4 °C to separate cell debris from the periplasmic fraction. The soluble fraction was then loaded onto a 5 mL HisTrap™ HP column (Cytiva) which had been equilibrated with binding buffer (20 mM Tris, pH 7.5, 150 mM NaCl, 30 mM imidazole). The column was then washed with 5 column volumes of binding buffer. Bound proteins were eluted using 5 column volumes of elution buffer (20 mM Tris, pH 7.5, 150 mM NaCl, 500 mM imidazole). The nanobodies were desalted using desalting buffer (20 mM Tris, pH 7.5, 150 mM NaCl) and further concentrated using a 3 kDa MWCO centrifugal filter unit (Amicon-Ultra 15). Nanobody solutions were aliquoted, and flash frozen in liquid nitrogen for long-term storage at –80 °C.

Expression of uniformly $^{15}N$-labelled nanobody mNb6-3C

Cells were grown at 37 °C until the OD reached 0.6–0.8 in Terrific Broth. Cells were centrifuged (5000 × *g*, 30 minutes, 4 °C) and resuspended in minimal medium (50 mM $NaHPO_4$, 25 mM

$KH_2PO_4$, 10 mM NaCl, pH 8.0) supplemented with 5 mM $MgSO_4$, 0.2 mM $CaCl_2$, 0.25% metal mix, $^{15}NH_4Cl$ (1g/L) as the only nitrogen source, and 1% glucose (*44*). Cells were grown in minimal medium at 37 °C for 30 minutes before induction using 1 mM IPTG and overexpression for 18–20 h at 25 °C. Cell lysis, extraction, and purification were performed as mentioned above. The $^{15}N$-labelled nanobody was buffer exchanged into NMR buffer (20 mM MES, pH 6.5, 150 mM NaCl, 10% $D_2O$), flash frozen in liquid nitrogen and stored at –80 °C.

<u>Expression and purification of mCherry</u>
The mCherry sequence (*24*) with C-terminal $His_6$ tag was cloned into the pET-29b(+) expression vector (Twist Bioscience, USA) and transformed into *E. coli* BL21 (DE3). The expression protocol was followed as described by Wang et al. with minor changes (*25*). Isolated single colonies were grown in 10 mL Luria Broth (LB) overnight, transferred to 1 L LB and grown at 37 °C until an OD of 0.8 was reached. This was followed by induction of overexpression using 0.2 mM IPTG for 18 h at 18 °C. The *E. coli* cells were harvested (5000 × *g*, 30 minutes, 4 °C) and resuspended in binding buffer (100 mM Tris, pH 8, 5% glycerol, 150 mM NaCl, 20 mM imidazole). Cells were lysed by sonication (Omni Sonic Ruptor 400 Ultrasonic homogenizer) three times at 50% power for 30 seconds on ice, before centrifugation at 17,000 × *g* for 30 minutes to separate cell debris. The soluble fraction was then loaded onto a 5 mL HisTrap™ HP column (Cytiva) which had been equilibrated with binding buffer. The column was then washed with 5 column volumes of the binding buffer. Protein was eluted using 5 column volumes of elution buffer (100 mM Tris, pH 8, 5% glycerol, 150 mM NaCl, 300 mM imidazole), desalted using desalting buffer (20 mM Tris, pH 7.5, 150 mM NaCl) and further concentrated using a 10 kDa MWCO centrifugal filter unit (Amicon-Ultra 15). The protein solution was aliquoted, and flash frozen in liquid nitrogen for long-term storage at –80 °C.

All proteins were characterized by SDS-PAGE and intact protein MS.

<u>Protein mass spectrometry (MS)</u>

<u>Intact protein MS</u>
Intact protein analysis was performed on Orbitrap Elite and Orbitrap Fusion™ Tribrid™ mass spectrometers (Thermo Fisher Scientific, USA) connected to a Thermo Fisher Scientific UltiMate 3000 HPLC system equipped with a ZORBAX 300SB-C3, 3.5 µm, 4.6 × 50 mm HPLC column (Agilent Technologies, USA). 10–20 µM protein samples were injected using a 500 µL/min linear gradient of solvent A (0.1% (*v*/*v*) formic acid in water) and solvent B (0.1% (*v*/*v*) formic acid in acetonitrile), ramping solvent B from 5% at the start to 80% after 12 min. The ion source was H-ESI with a static spray voltage set to 3500 V in positive ion mode. The sheath, auxiliary and sweep gasses were set to 50 Arb, 10 Arb and 1 Arb, respectively. The ion transfer tube temperature was set to 325 °C and the vaporizer temperature was set to 350 °C. The resolution of the Orbitrap MS detector was set to 120000, and the scan range was set to 500–2000 m/z. Protein mass was determined by deconvolution using the program Xcalibur 3.0.63 (Thermo Fisher Scientific, USA).

<u>Native MS with prior buffer exchange</u>
Samples for native MS were buffer exchanged into 100 mM $NH_4OAc$ (pH 7) in a 3 kDa MWCO centrifugal filter unit (Amicon-Ultra 0.5) prior to analysis. Native protein analysis was performed on an Orbitrap Fusion™ Tribrid™ mass spectrometer (Thermo Fisher Scientific, USA) connected to a Thermo Fisher Scientific UltiMate 3000 HPLC system. 10–20 µM protein samples were injected using a 5 minute 100 µL/min isocratic elution mode in 100 mM $NH_4OAc$ (pH 7). The ion

source was H-ESI with a static spray voltage set to 3000 V in positive ion mode. The sheath, auxiliary and sweep gasses were set to 25 Arb, 5 Arb and 0 Arb, respectively. The ion transfer tube temperature was set to 275 °C and the vaporizer temperature was set to 50 °C. The MS detector was an Orbitrap with the resolution set to 120000, and the scan range was set to 500–4000 m/z. Native protein mass was determined by deconvolution using the program Xcalibur 3.0.63 (Thermo Fisher Scientific, USA).

Native MS with online buffer exchange

Native protein analysis was performed on an Orbitrap Fusion™ Tribrid™ mass spectrometer (Thermo Fisher Scientific, USA) connected to a Thermo Fisher Scientific UltiMate 3000 HPLC system equipped with a NativePac OBE-1, 3 µm, 2.1 × 50 mm HPLC column (Thermo Scientific, Australia). 10–50 µM protein samples were injected using a 3 minute 100 µL/min isocratic elution mode in 100 mM $NH_4OAc$ (pH 7). The divert valve was set to send the flow to the source for the first 1.1 minutes before sending the flow to the waste for the remainder of the run. The ion source was H-ESI with a time dependent spray voltage set to 3000 V in positive ion mode for the first 2 minutes before changing to 0 for the remainder of the run. The sheath, auxiliary and sweep gasses were set to 25 Arb, 5 Arb and 0 Arb, respectively. The ion transfer tube temperature was set to 275 °C and the vaporizer temperature was set to 50 °C. The MS detector an Orbitrap with the resolution set to 120000, and the scan range set to 500–4000 m/z. Native protein mass was determined by deconvolution using the program Xcalibur 3.0.63 (Thermo Fisher Scientific, USA).

Metal uptake reaction optimization

Initial attempts to incorporate Bi(III) into nanobodies involved incubating mNb6-3C in 1 M and 6 M guanidinium hydrochloride (GdmCl) for 20 minutes at room temperature (RT), using 25 mM TCEP at pH 3. These conditions resulted in low metal uptake, and the removal of excess denaturant (GdmCl) to obtain functional protein proved challenging. Consequently, further optimizations were focused on applying simultaneous temperature and pH shock at different time points.

For Bi(III), In(III) and Ga(III) binding optimization, the nanobodies (50–100 µM) were reduced either at RT or at a temperature range 8–10 °C below their respective denaturation midpoint ($T_m$) in absence (pH 7.5) or presence of 1 (pH 6.5), 5 (pH 4.0), 10 (pH 3.5) or 25 (pH 3) mM TCEP for a maximum of 120 minutes. Reduced nanobodies were aliquoted at different time points, followed by the addition of 1–5 equivalents of the metal salts $BiBr_3$ and gastrodenol (for bismuth), $InCl_3$ hexahydrate (for indium), and $Ga(NO_3)_3$ hydrate (for gallium) to the reaction solution from their respective stocks, and immediately vortexed for 60 s. This protocol was effective for the mNb6 and 2Rs15d nanobody libraries. For the Lam2 nanobody subset, gastrodenol (for Bi(III) uptake) was co-incubated with the nanobodies for up to 120 minutes. Following the reaction, the solution was quickly centrifuged, and the supernatant was collected for further analysis. For quantifying metal uptake of mNb6-3C, mNb6-3C* and mNb6-4C by native MS, the supernatant was buffer exchanged to 100 mM $NH_4OAc$ (pH 7). For native MS of Lam2-3C-Bi using an online buffer exchange (OBE) column, the supernatant was directly used for data acquisition and analyses. Detailed optimization conditions for the metal-uptake reactions of the individual nanobodies are outlined in Table S1.

**Table S1.** Conditions tested during optimization for metal uptake (RT = room temperature).

| Nanobody | Temperature (°C) | [TCEP] mM | pH | Time (min) | Eq. of metal salts |
|---|---|---|---|---|---|
| mNb6-3C | RT, 50, 55, 60, 70 | 1, 5, 10, 25 | 7.5, 6.5, 4.0, 3.5, 3 | up to 90 | 1–5 |
| mNb6-3C* | 50 | 25 | 3 | up to 60 | 5 |
| mNb6-4C | 50, 55, 60 | 25 | 3 | up to 90 | 5 |
| Lam2-3C | RT, 58 | 25 | 3 | up to 120 | 5 |
| 2Rs15d-5C | 37, 50 | 25 | 3 | up to 60 | 5 |
| 2Rs15d-2A-3C | 50 | 25 | 3 | up to 60 | 5 |

Conditions which resulted in >98% metal uptake (Table S2), without denaturing the nanobodies were subsequently used for all other experiment and assays.

**Table S2.** Optimized conditions for metal uptake.

| Nanobody | Temperature (°C) | [TCEP] mM | pH | Time (min) | Eq. of metal salts |
|---|---|---|---|---|---|
| mNb6-3C | 50 | 25 | 3.0 | 5, 15 | 5 |
| mNb6-3C* | 50 | 25 | 3.0 | 30 | 5 |
| mNb6-4C | 60 | 25 | 3.0 | 15 | 5 |
| Lam2-3C | 58 | 25 | 3.0 | 60 | 5 |
| 2Rs15d-5C | 37 | 25 | 3.0 | 30 | 5 |
| 2Rs15d-2A-3C | 50 | 25 | 3.0 | 30 | 5 |

<u>Evaluating the stability of metal-bound nanobodies</u>
Nanobodies were reduced and saturated with either Bi(III), In(III) or Ga(III) as per the optimized protocol to obtain >98% metal saturation, prior to buffer exchange into 100 mM $NH_4OAc$ (pH 7). Samples were then stored at 4 °C for up to 14 days and analyzed *via* native MS at different time points to determine the stability of the metal-bound nanobody.

<u>Assessing metal uptake of ‘primed’ (pre-reduced) nanobodies</u>
Nanobodies were reduced as described above and stored at 4 °C for up to 14 days prior to the addition of either Bi(III), In(III) or Ga(III) as per the optimized protocol to obtain >98% metal saturation. At different time points, metal-bound nanobodies were then buffer exchanged into 100 mM $NH_4OAc$ (pH 7) and analyzed *via* native MS to determine the ability of the pre-reduced (‘primed’) nanobodies to bind to metals.

<u>Dialysis to assess stability of metal nanobodies</u>
In order to monitor any potential metal dissociation and rule out the impact of excess metal salts and TCEP on the stability of the nanobodies, the metal-bound nanobodies were dialysed (1000:1 *v*/*v*) overnight for 18 h at 4 °C in 20 mM HEPES, pH 7.8, 150 mM NaCl. mNb6-4C-In and mNb6-4C-Ga were co-dialysed in separate 3 kDa MWCO membranes within the same dialysis tank under the above conditions to identify metal cross-contamination. Dialysed nanobodies were then examined *via* native MS.

Glutathione (GSH) competition assay

At first, reduction followed by uptake reactions with Bi(III), In(III) and Ga(III) were performed with the nanobodies (mNb6-3C, mNb6-3C* and mNb6-4C) as per the optimized protocol to obtain >98% metal saturation. The metal-bound nanobody samples were buffer exchanged to 100mM $NH_4OAc$ (pH 7). A reduced glutathione (GSH) stock was prepared in 100 mM $NH_4OAc$ and adjusted to pH 7. Competition experiments were performed with 1, 10, 20, 50, and 100 equiv. of GSH with respect to the concentrations of the metal-bound nanobodies (50 μM) for 1 h at 25 °C. Native MS of the nanobodies was performed in 100 mM $NH_4OAc$. Metal uptake in nanobodies (%) was plotted as a function of increasing equivalents of GSH in GraphPad Prism 10 (Dotmatics, USA). One-phase exponential decay was used as the fitting function.

Competition with human apo-transferrin

Similar to the GSH competition assay protocol, Bi(III), In(III) and Ga(III) uptake reactions were performed with the nanobodies (mNb6-3C, mNb6-3C* and mNb6-4C) as per the optimized protocol to obtain >98% metal saturation. Human apo-transferrin from Sigma-Aldrich (T4382) was resuspended in 100 mM $NH_4OAc$ (pH 7) to prepare a stock concentration of 500 μM. Competition experiments were performed at 1, 2, and 4 equiv. of apo-transferrin with respect to the concentrations of the metal-bound Nbs (25 μM) for 1 h at 25 °C. Native MS of the nanobodies was performed in 100 mM $NH_4OAc$. Metal uptake in nanobodies (%) was plotted as a function of increasing equivalents of apo-transferrin in GraphPad Prism 10 (Dotmatics, USA). One-phase exponential decay was used as the fitting function.

Circular dichroism (CD) spectroscopy

Secondary structures of the nanobodies were assessed by circular dichroism using a Chirascan spectropolarimeter from Applied Photophysics equipped with a temperature control module using a 0.1 cm path-length cuvette. Each CD spectrum was averaged over 2 scans, and the baseline correction was done by subtraction of the spectrum with the appropriate blank solution. Individual nanobodies were diluted to 10–20 μM in 20 mM phosphate buffer, pH 7.5. Scans were carried out at 25 °C over a range of 200–260 nm with a 0.5 nm step size and 1 nm bandwidth. Thermal denaturation experiments were monitored at 204 nm (mNb6, Lam2 nanobodies) or 222 nm (2Rs15d nanobodies) over a temperature range of 20–80 °C (1 °C/min heating rate). Temperature-dependent CD data were fitted to a two-state unfolding model (Boltzmann sigmoidal equation) and plotted in GraphPad Prism 10 (Dotmatics, USA) to obtain the denaturation midpoint ($T_m$).

NMR spectroscopy

All NMR spectra were recorded at 25 °C using an 800 MHz Bruker Avance NMR spectrometer equipped with a cryoprobe. [$^{15}N$, $^{1}H$]-HSQC spectra were recorded in a 3 mm NMR tube (sample volume 180 μL). The NMR spectrum of mNb6-3C was acquired in NMR buffer (20 mM MES, pH 6.5, 150 mM NaCl, 10% $D_2O$). For Bi(III) uptake, the protein sample was subjected to heating at 50 °C in presence of 10 mM TCEP (pH 6.5) for 15 minutes. 5 equiv. of gastrodenol (bismuth source) were added and the sample was immediately vortexed. After centrifugation, the sample concentration was 0.15 mM. The [$^{15}N$, $^{1}H$]-HSQC spectrum of mNb6-3C-Bi was acquired with the same parameters as for the apo-nanobody.

Isothermal titration calorimetry (ITC)

mCherry, Lam2, and Lam2-3C-Bi were dialyzed overnight at 4 °C in 20 mM HEPES, pH 8, 150 mM NaCl (ITC buffer) in 10 MWCO (mCh) and 3 MWCO (Nb) dialysis tubings from SpectraPor®. Post-dialysis, proteins were concentrated using Amicon-Ultra 15 centrifugal filter units with 10 MWCO (mCh) and 3 MWCO (Nbs) to 700 μM and 100 μM, respectively.

Titrations were performed on a TA Instruments Benchtop Nano ITC. All stocks and dilutions were prepared using the buffer from the overnight dialysis. mCherry and the corresponding nanobody samples were degassed prior to each ITC experiment. All titrations were performed as titrations of mCherry (protein) into nanobodies (ligands) at 25 °C with a stirring rate of 350 rpm (conditions outlined in Table S3). Initial and final baselines were generated over 120 s. The first injection of each titration was a 1 μL blank injection, followed by 22 injections of 2 μL each, with an injection interval of 300 s. The background heat was estimated as the average heat associated with each injection in a control titration of protein into buffer and subtracted from each titration. Results were analyzed in NITPIC for baseline detection, blank subtraction, and integration of baseline-subtracted power (*45*). SEDPHAT was used to fit integrated heats to the single binding site model (A + B ←→ AB, hetero association) by global fitting with the Simplex algorithm (*46*). Thermograms were plotted in GUSSI (*47*).

**Table S3.** Sample details for ITC experiments.

| Interaction | [Nanobody] (Syringe) | [mCherry] (Cell) | Molar ratio (S:C) |
|---|---|---|---|
| Lam2 & mCherry | 670 μM | 67 μM | 10:1 |
| Lam2-3C-Bi & mCherry | 200 μM | 40 μM | 5:1 |

Surface plasmon resonance (SPR)

SPR measurements were performed with 20 mM HEPES, pH 7.8, 150 mM NaCl, 0.05% Tween-20 as running buffer. For each experiment, 100–150 μg of metal nanobodies (>98% metal saturation confirmed by native MS), mNb6-3C-Bi, mNb6-4C-In and mNb6-4C-Ga, in 10 mM sodium acetate buffer, pH 4.8 were freshly immobilised on a CM5, Series S sensor chip flow-cell (Cytiva, 29104988, USA) using EDC/NHS chemistry at 25 °C. The contact time for ligand binding was set to 1800 s with a flow rate of 5 μl/min. The unbound sites were blocked using ethanolamine hydrochloride solution. An amine coupling kit (Cytiva, BR100050, USA) was used for the coupling step. For the experiments, the response unit (RU) after coupling ranged from 300–1000. The initial concentration of the analyte, SARS-CoV-2 receptor binding domain (RBD), ranged from 25 to 50 nM and was serially diluted down to 0.19 nM to evaluate its binding affinity to the nanobodies. All binding experiments were conducted at 20 °C in single-cycle kinetics mode. SPR experiments were performed on a Biacore 8K (Cytiva, USA). The instrument was operated using Biacore Insight 5 software (Cytiva, USA). The data were plotted and analyzed in GraphPad Prism 10 (Dotmatics, USA) using the non-linear fit 'one site – specific binding'. $K_D$ values of duplicate measurements are represented as the mean value ± standard deviation.

Native gel electrophoresis

The Lam2-3C-Bi:mCherry complex was created by incubating the metal-bound nanobody (>98% metal saturation confirmed by native MS) and mCherry together (1:1 ratio) in 20 mM Tris, pH 7.5, 150 mM NaCl on ice for 60 minutes. The NativePAGE Novex Bis-Tris Gel system protocol was followed with minor changes. 15 μL of metal-bound nanobody:protein complexes were added to 5 uL of 4 × loading dye (200 mM BisTris-HCl, 200 mM NaCl, 40% glycerol, 0.004% Ponceau S, pH 7.2). The gel was washed 3 times with running buffer (50 mM BisTris, 50 mM tricine, pH 6.8) and 20 μL of sample was loaded. The gel was run in running buffer at 4 $^{o}$C for 60 min at 150 V, then for further 45 min at 250 V (Thermo EC250-90 powerpack). The gel was viewed under UV light on a Biorad ChemiDock MP imaging system to observe fluorescent proteins, then fixed in 40% methanol, 10% acetic acid for 30 min and stained with 0.02% Coomassie R-250 in 30% methanol and 10% acetic acid for 30 min before destaining in 8% acetic acid until the desired background was obtained. The gel was visualized with a Biorad ChemiDock MP imaging system.

SDS PAGE with chemical cross linking

The mNb6-3C-Bi:SARS-CoV-2 receptor binding domain (RBD) complexes were created by incubating the metal-bound nanobody (>98% metal saturation confirmed by native MS) and SARS-CoV-2 RBD together in phosphate buffered saline (PBS) on ice for 60 minutes. Nanobody:protein complexes were crosslinked by incubation with 100 equiv. (relative to nanobody:protein concentration) of disuccinimidyl glutarate on ice for 60 minutes. The samples were then prepared and run on SDS PAGE following standard running and staining procedures. The gel was visualized with a Biorad ChemiDock MP imaging system.

Inductively Coupled Plasma Mass Spectrometry (ICP-MS)

Relevant lanes of the native PAGE gel were excised with a scalpel and soaked in concentrated nitric acid for 10 minutes. The samples were then diluted to 10% nitric acid for analysis. A ThermoFisher iCap RQ ICP-MS was used to measure bismuth abundance in the samples (measured in ppb). A calibration curve (0.1 ppb, 1 ppb, 10 ppb, and 100 ppb) of Agilent Intelliquant standard with the isotope $^{209}$Bi was run prior to the diluted gel samples, with calibration curve fits of better than 0.999 accepted.

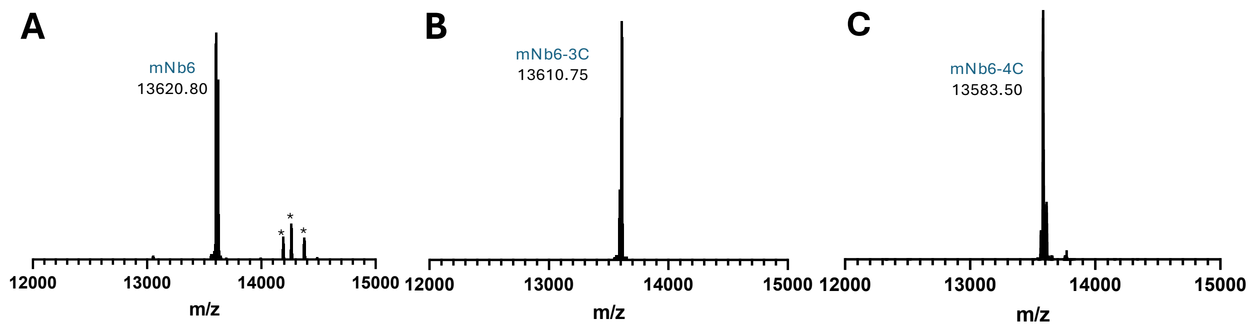


**Fig. S1.** Intact MS of mNb6 (A), mNb6-3C (B), and mNb6-4C (C) after time-dependent reduction and modification with 25 mM TCEP and 5 equiv. gastrodenol at room temperature. The deconvoluted m/z ratios indicate the presence of an intact disulfide bond. Species of higher m/z ratio that correspond to a partially cleaved N-terminal leader sequence are indicated with an asterisk (*).

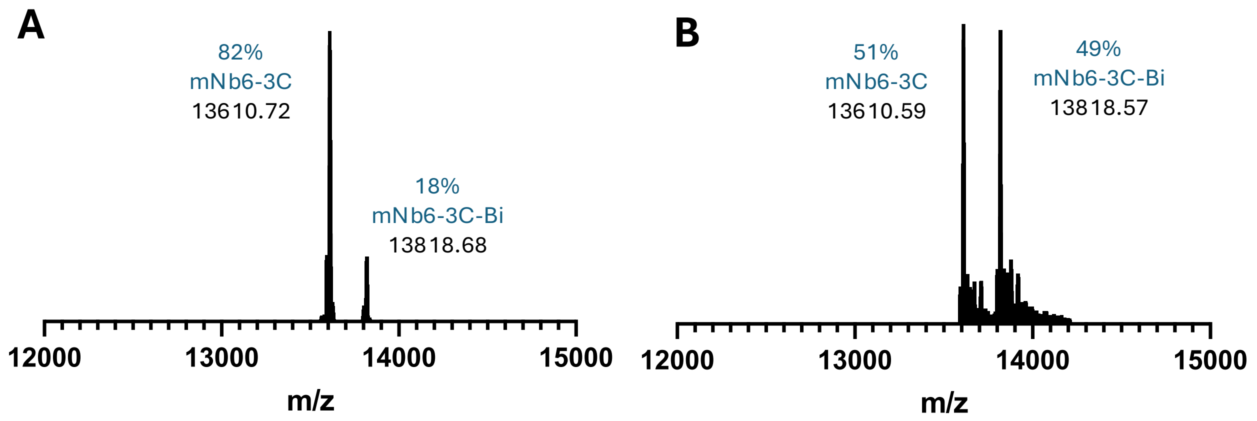


**Fig. S2.** Native MS of mNb6-3C after treatment with 1 M (A) and 6 M (B) guanidinium hydrochloride at room temperature for 20 min in presence of 25 mM TCEP (pH 3), followed by addition of 5 equiv. $BiBr_3$.

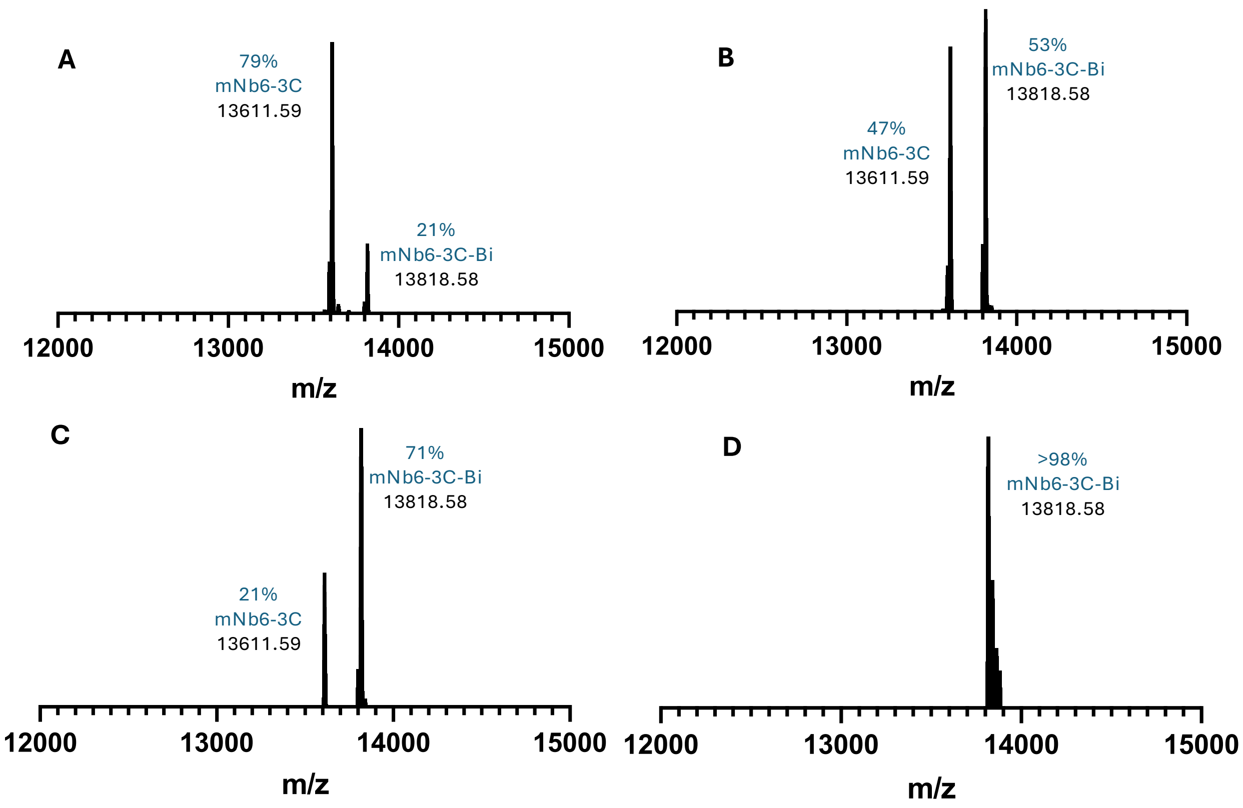


**Fig S3.** Representative native MS showing quantitative Bi(III) uptake (from 5 equiv. gastrodenol) by mNb6-3C after incubation with 1 mM TCEP, pH 6.5 (A), 5 mM TCEP, pH 4 (B), 10 mM TCEP, pH 3.5 (C) and 25 mM TCEP, pH 3 (D) for 15 min at 50 °C.

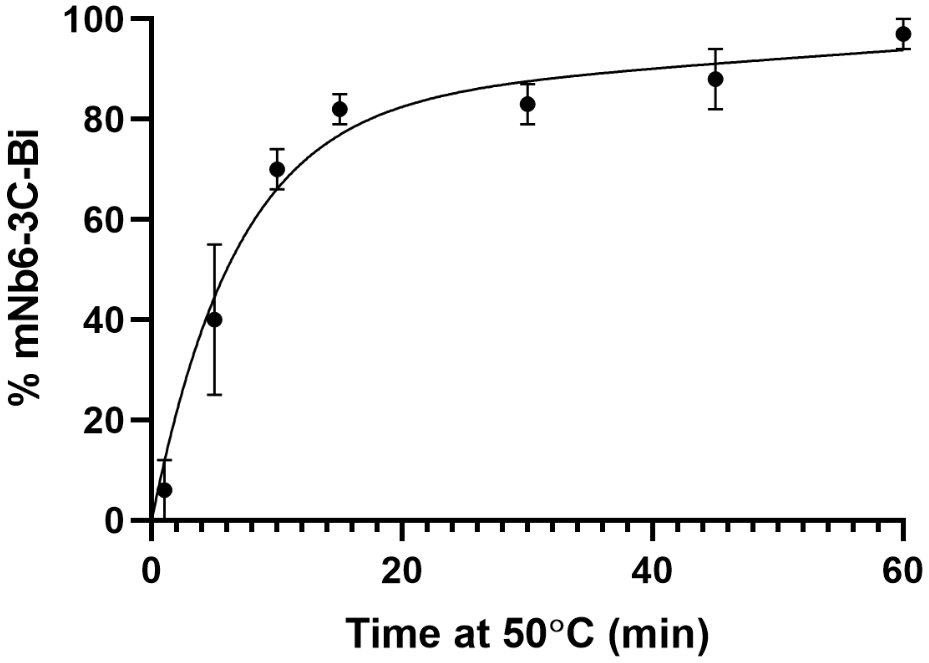


**Fig. S4.** Uptake of Bi(III) by mNb6-3C from $BiBr_3$ (DMSO stock) after reduction with 25 mM TCEP at pH 3 and 50 °C heat shock over different durations.

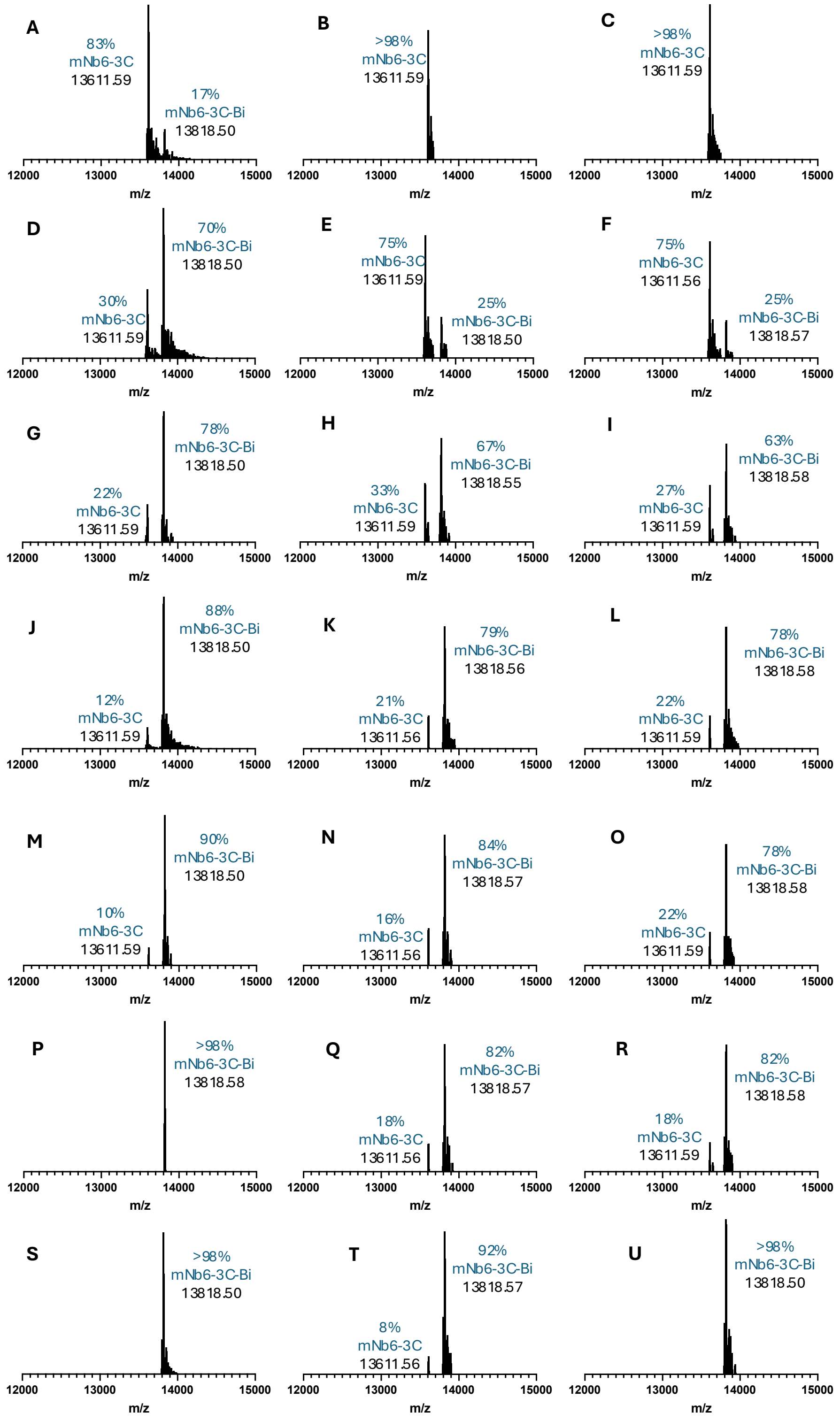


**Fig. S5.** Optimization of Bi(III) uptake by mNb6-3C using $BiBr_3$. Native mass spectra of samples measured in triplicate after 1 min (A to C), 5 min (D to F), 10 min (G to I), 15 min (J to L), 30 min (M to O), 45 min (P to R), and 60 min (S to U) incubation at 50 °C in 25 mM TCEP at pH 3.0, followed by addition of $BiBr_3$ (DMSO stock).

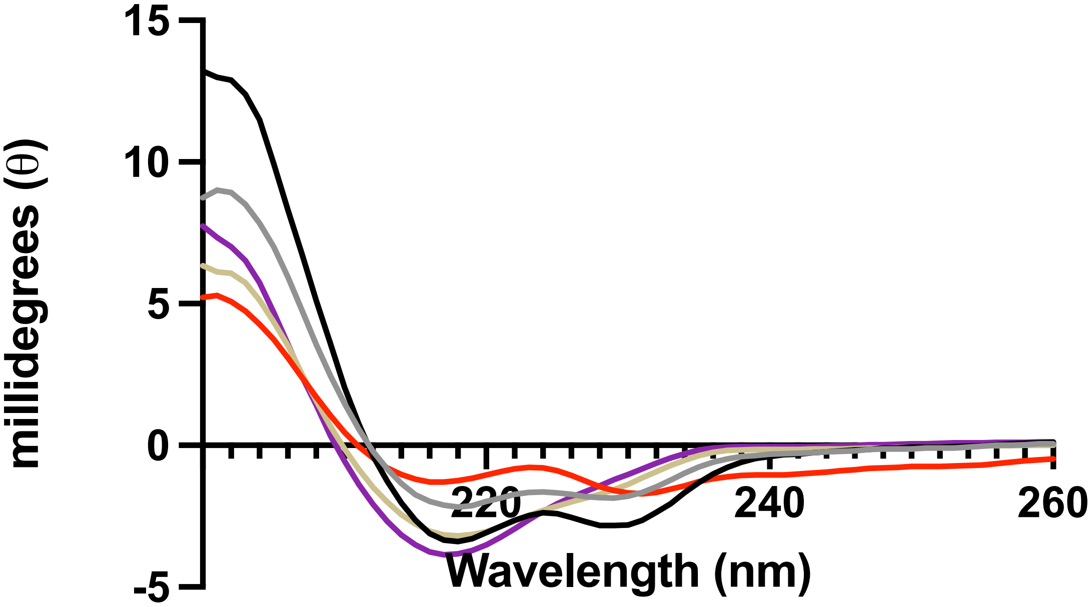


**Fig. S6.** Circular dichroism (CD) spectra of mNb6 (grey), mNb6-3C (black), mNb6-3C-Bi (red), mNb6-3C* (pale yellow), and mNb6-3C*-Bi (purple) measured at 15 $\mu$M in 20 mM phosphate buffer, pH 7.5.

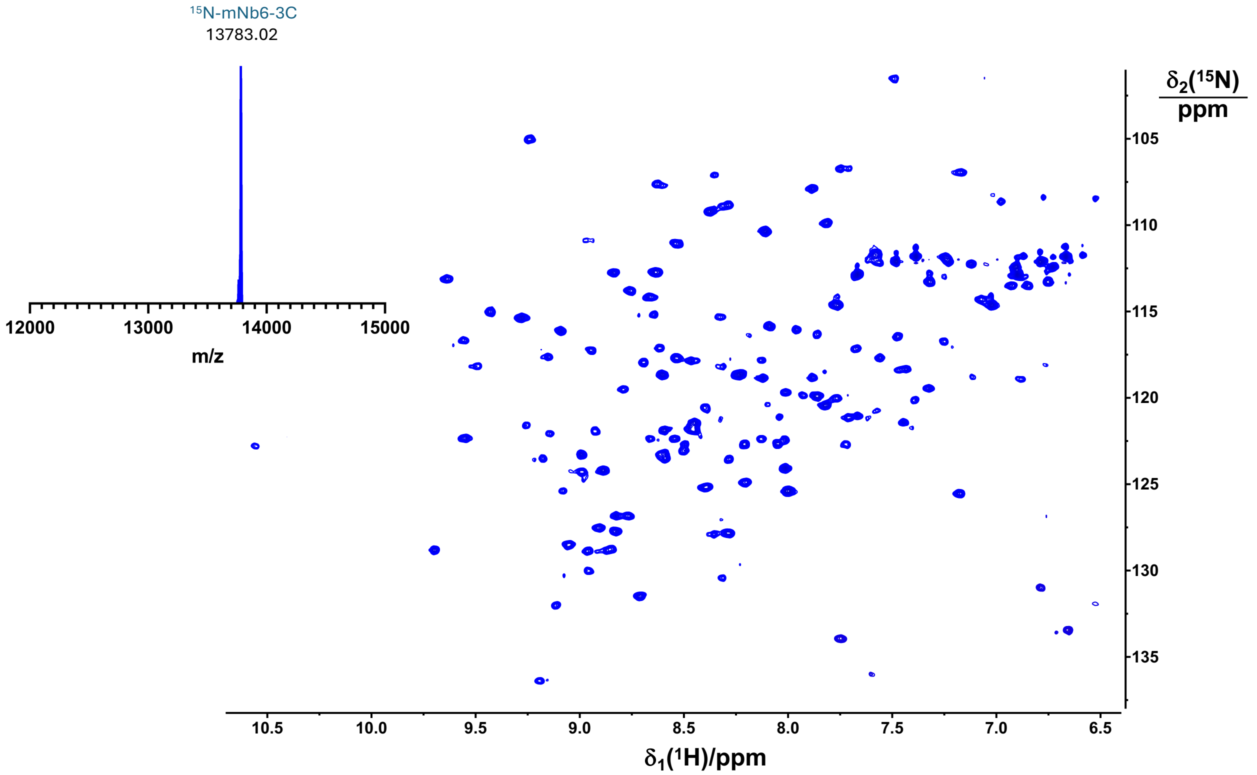


**Fig. S7.** 800 MHz [$^{15}N$,$^{1}H$]-HSQC NMR spectrum of a 150 µM solution of $^{15}N$-mNb6-3C in 20 mM MES pH 7.5, 150 mM NaCl, 10% $D_2O$. Intact MS spectrum of the protein is shown in the inset.

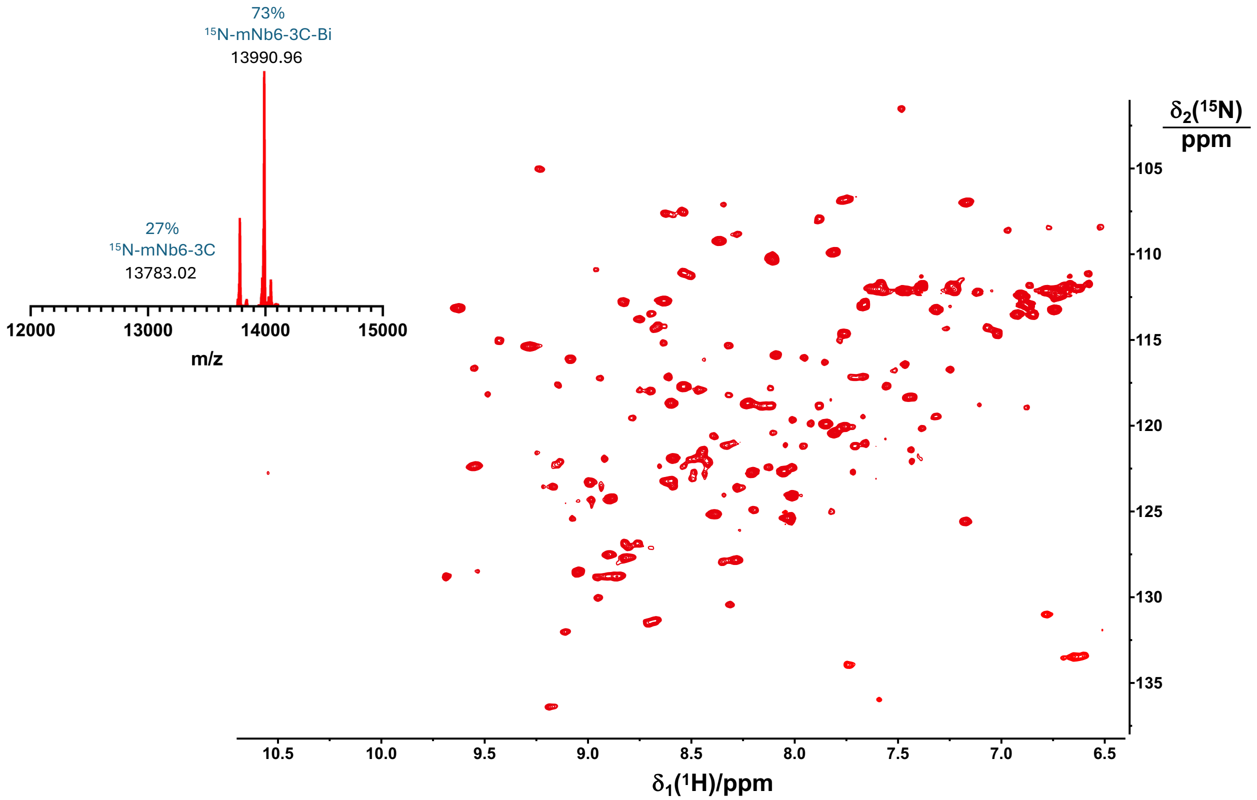


**Fig. S8.** 800 MHz [$^{15}N$,$^{1}H$]-HSQC NMR spectrum of a 150 µM solution of $^{15}N$-mNb6-3C-Bi in 20 mM MES pH 7.5, 150 mM NaCl, 10 mM TCEP, 10% $D_2O$. Native MS spectrum of the protein is shown in the inset.

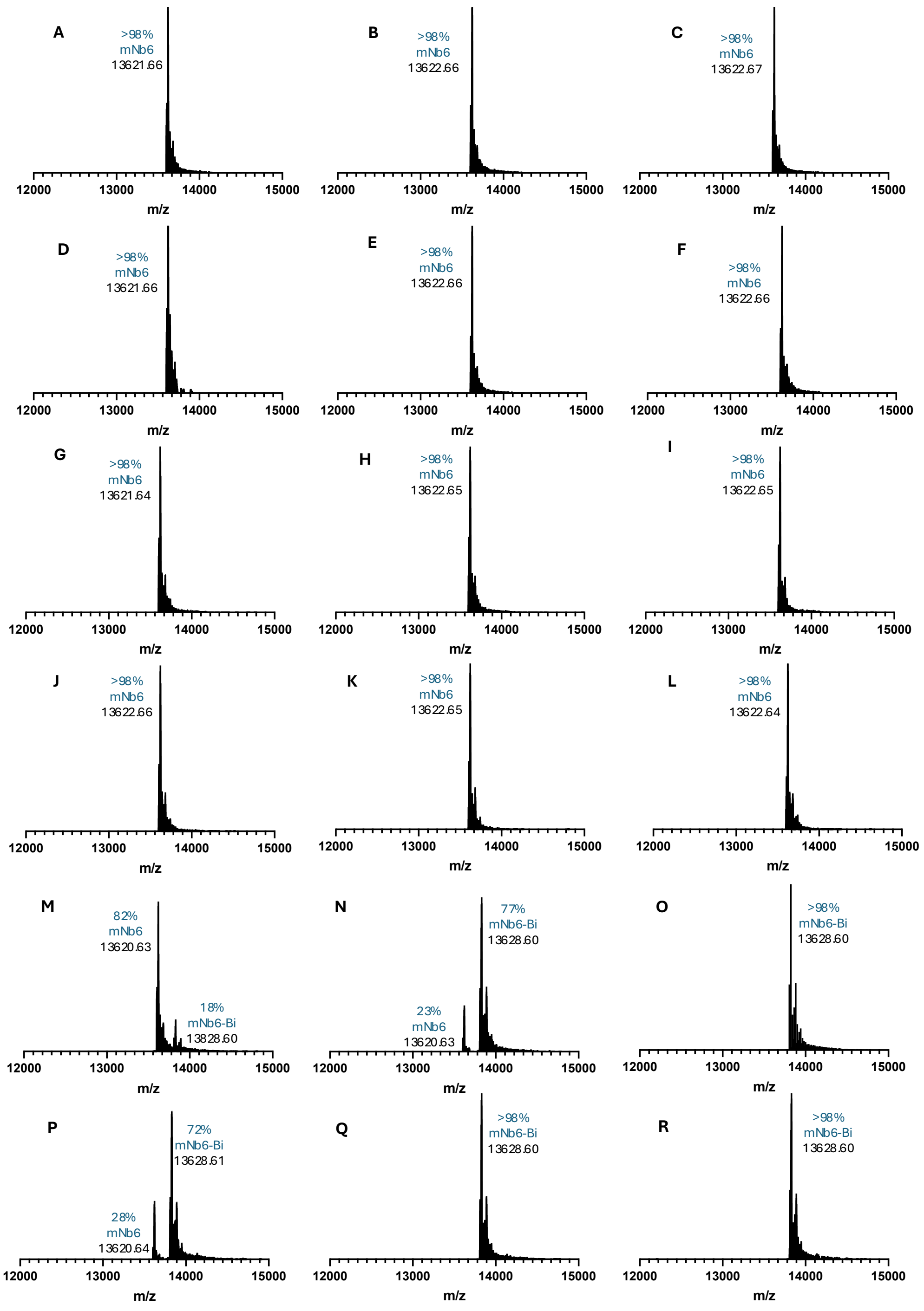


**Fig. S9.** Uptake of In(III), Ga(III) and Bi(III) by mNb6 determined using native MS after incubation at specified temperature and time in 25 mM TCEP at pH 3. (A) In(III), 50 °C, 15 min; (B) In(III), 55 °C, 15 min; (C) In(III), 60 °C, 15 min; (D) In(III), 50 °C, 90 min; (E) In(III), 55 °C, 90 min; (F) In(III), 60 °C, 90 min; (G) Ga(III), 50 °C, 15 min; (H) Ga(III), 55 °C, 15 min; (I) Ga(III), 60 °C, 15 min; (J) Ga(III), 50 °C, 90 min; (K) Ga(III), 55 °C, 90 min; (L) Ga(III), 60 °C, 90 min; (M) Bi(III), 50 °C, 15 min; (N) Bi(III), 55 °C, 15 min; (O) Bi(III), 60 °C, 15 min; (P) Bi(III), 50 °C, 90 min; (Q) Bi(III), 55 °C, 90 min; (R) Bi(III), 90 °C 90 min.

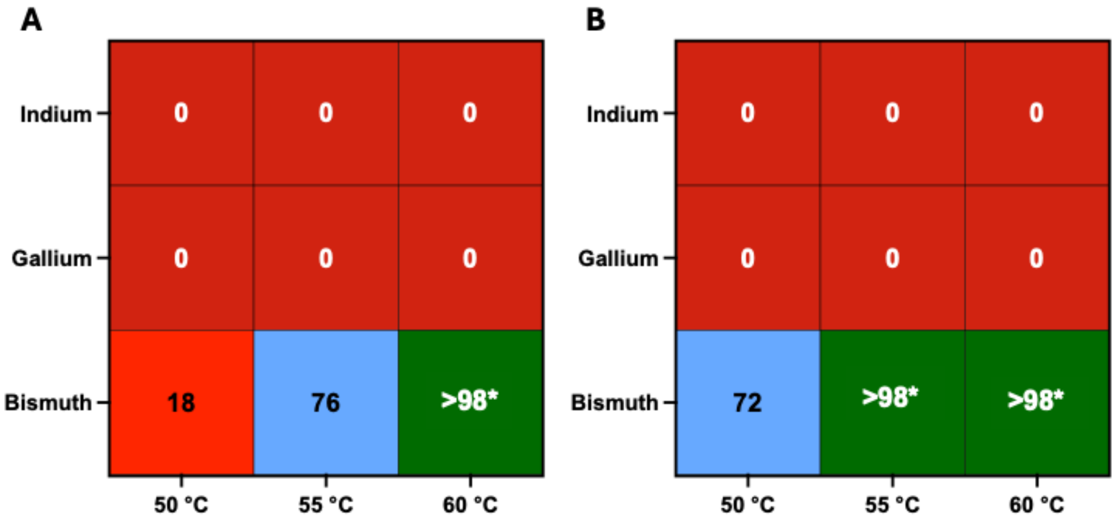


**Fig. S10.** Heatmap of In(III), Ga(III) and Bi(III) uptake by mNb6 following incubation at indicated temperatures for 15 (A) and 90 minutes (B). Uptake determined by native MS is indicated in %. Conditions where excessive protein precipitation was observed are indicated by an asterisk (*).

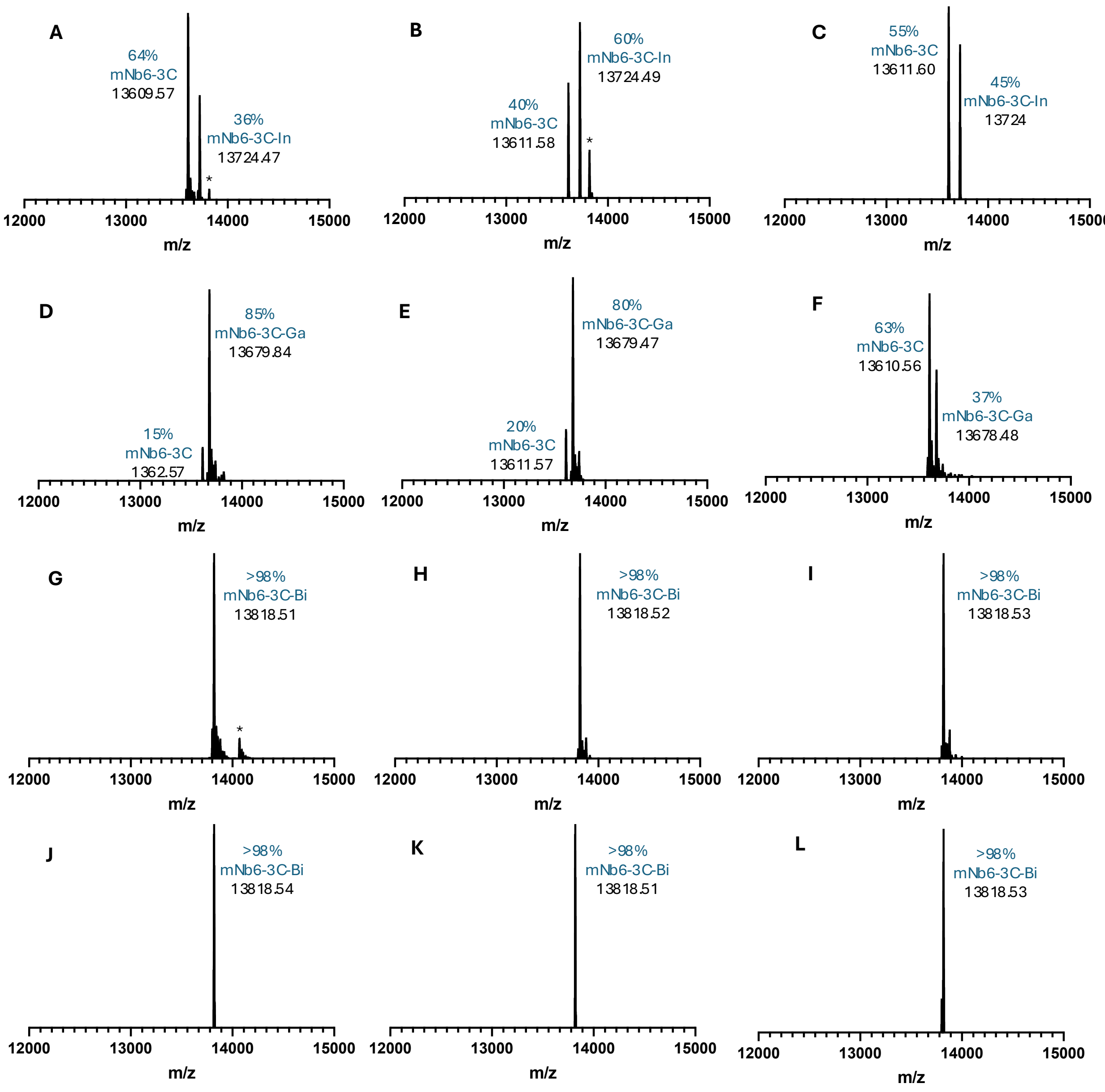


**Fig. S11.** Uptake of In(III), Ga(III) and Bi(III) by mNb6-3C determined using native MS after incubation at specified temperature and time in 25 mM TCEP at pH 3. (A) In(III), 50 °C, 15 min; (B) In(III), 55 °C, 15 min; (C) In(III), 60 °C, 15 min; (D) Ga(III), 50 °C, 15 min; (E) Ga(III), 55 °C, 15 min; (F) Ga(III), 60 °C, 15 min; (G) Bi(III), 50 °C, 15 min; (H) Bi(III), 55 °C, 15 min; (I) Bi(III), 60 °C, 15 min; (J) Bi(III), 50 °C, 90 min; (K) Bi(III), 55 °C, 90 min; (L) Bi(III), 60 °C, 90 min. The asterisks (*) indicate unknown protein impurities.

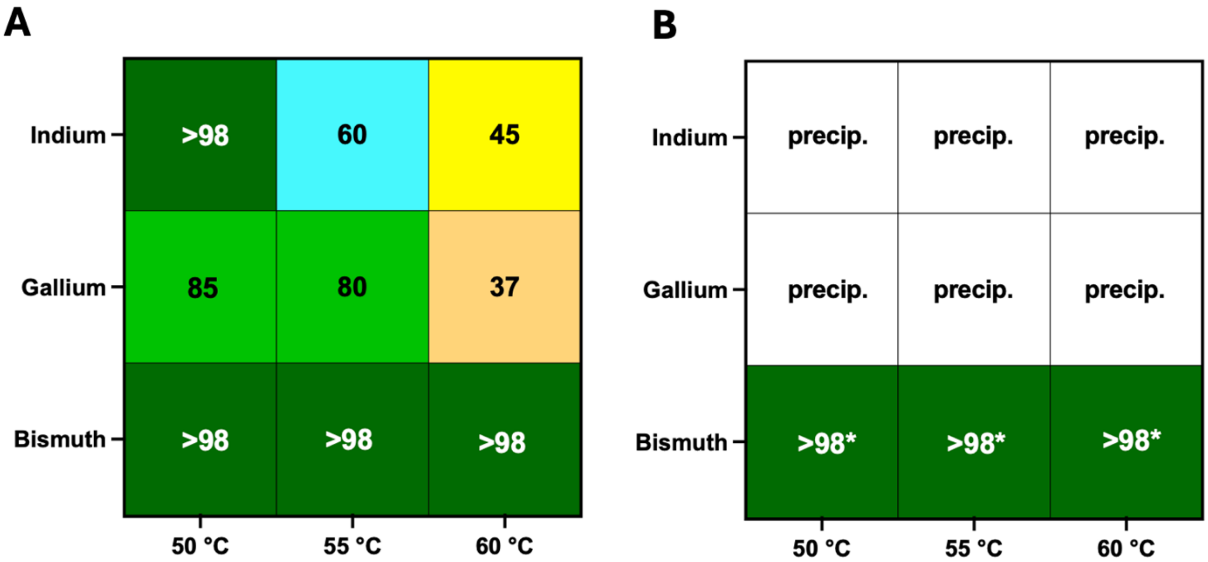


**Fig. S12.** Heatmap of In(III), Ga(III) and Bi(III) uptake by mNb6-3C following incubation at indicated temperatures for 15 (A) and 90 minutes (B). Uptake determined by native MS is indicated in %. Conditions where excessive precipitation was observed are indicated by an asterisk (*).

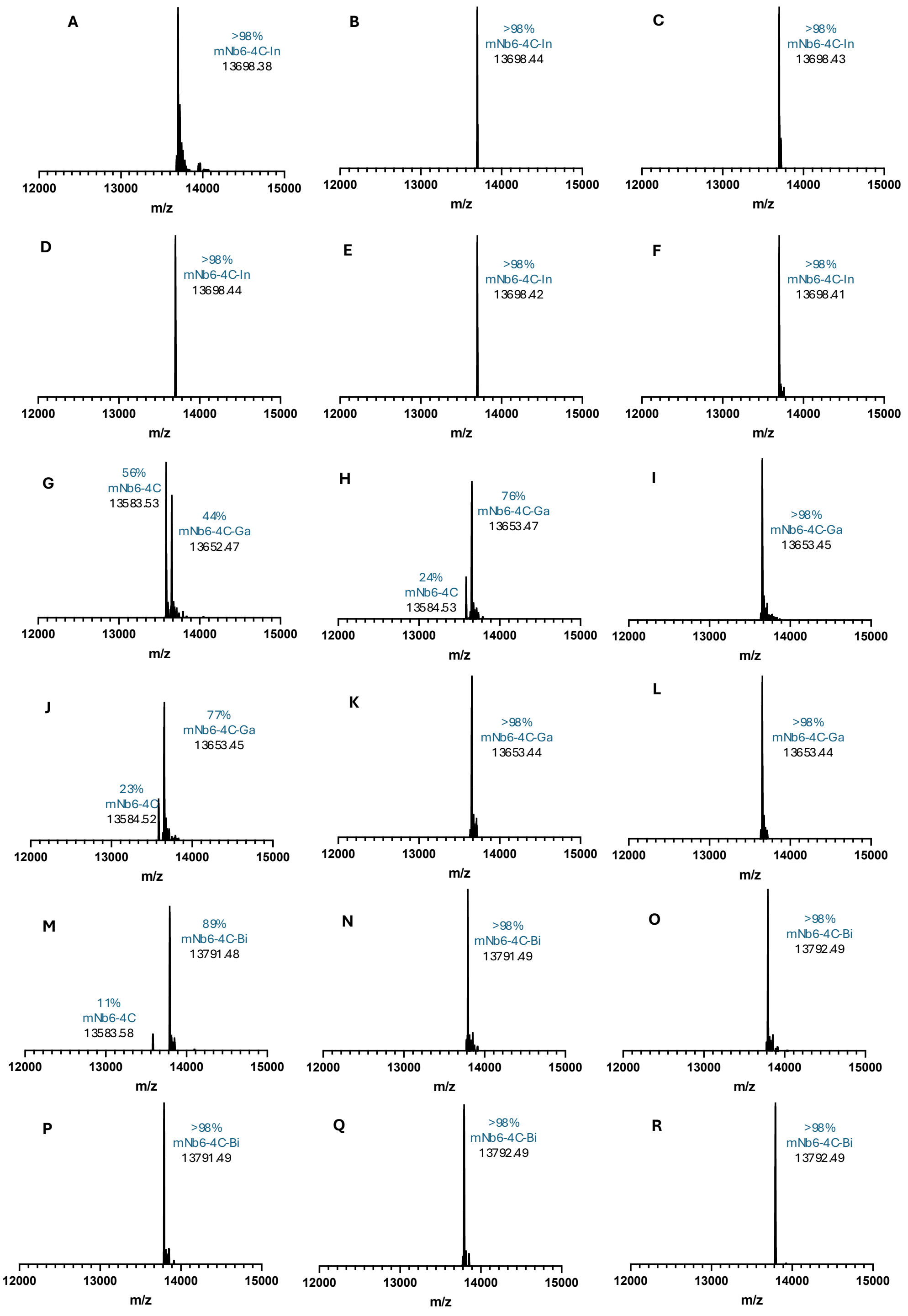


**Fig. S13.** Uptake of In(III), Ga(III) and Bi(III) by mNb6-4C determined using native MS after incubation at specified temperature and time in 25 mM TCEP at pH 3. (A) In(III), 50 °C, 15 min; (B) In(III), 55 °C, 15 min; (C) In(III), 60 °C, 15 min; (D) In(III), 50 °C, 90 min; (E) In(III), 55 °C, 90 min; (F) In(III), 60 °C, 90 min; (G) Ga(III), 50 °C, 15 min; (H) Ga(III), 55 °C, 15 min; (I) Ga(III), 60 °C, 15 min; (J) Ga(III), 50 °C, 90 min; (K) Ga(III), 55 °C, 90 min; (L) Ga(III), 60 °C, 90 min; (M) Bi(III), 50 °C, 15 min; (N) Bi(III), 55 °C, 15 min; (O) Bi(III), 60 °C, 15 min; (P) Bi(III), 50 °C, 90 min; (Q) Bi(III), 55 °C, 90 min; (R) Bi(III), 90 °C 90 min.

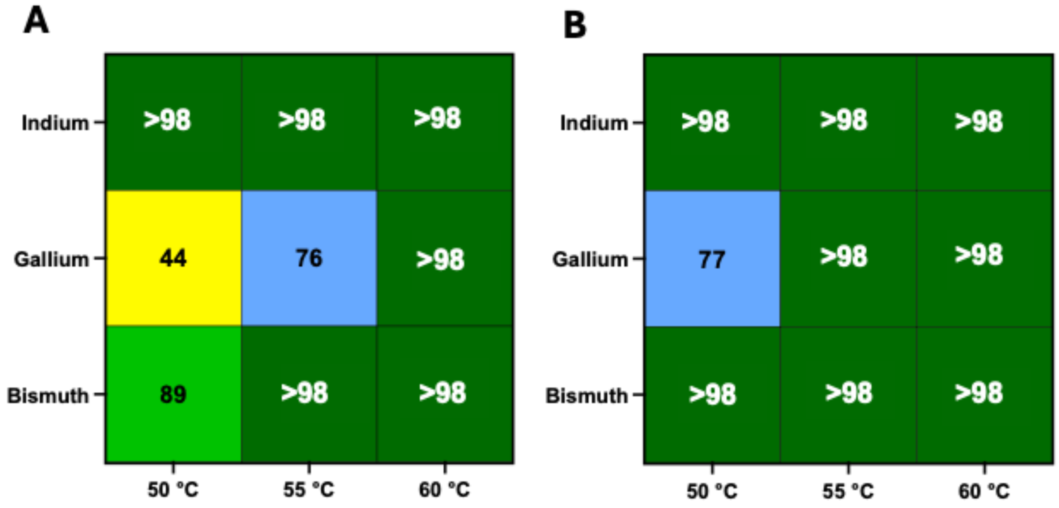


**Fig. S14.** Heatmap of In(III), Ga(III) and Bi(III) uptake by mNb6-4C following incubation at indicated temperatures for 15 (A) and 90 minutes (B). Uptake determined by native MS is indicated in %.

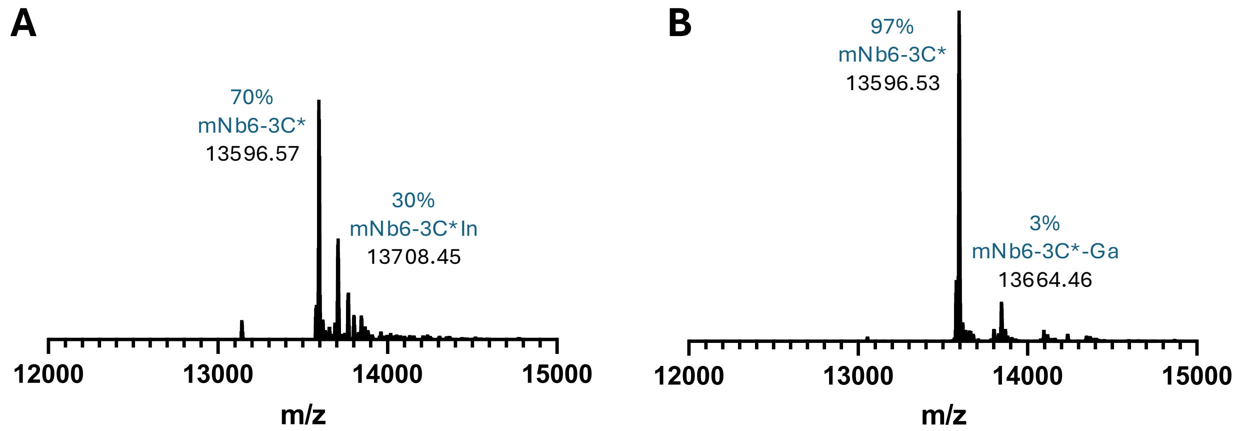


**Fig. S15.** Native MS of mNb6-3C* after incubation at 50 ºC in 25 mM TCEP, pH 3.0 for 15 minutes, followed by addition of In(III) (A) and Ga(III) (B).

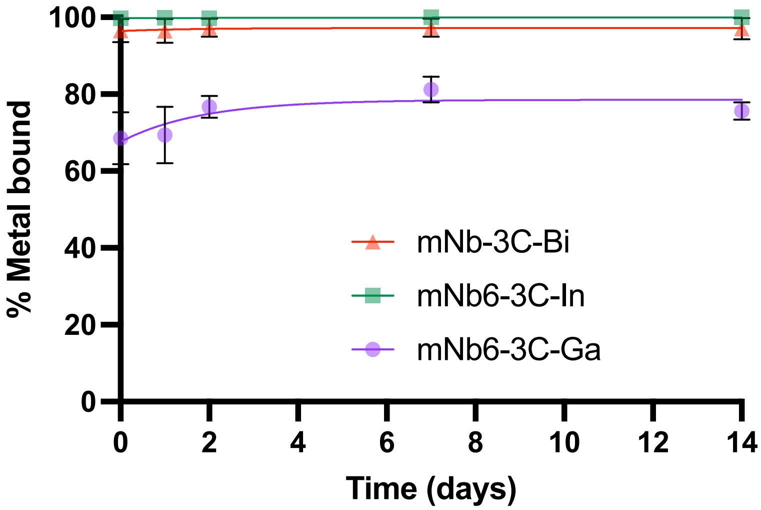


**Fig S16.** Stability of mNb6-3C in complex with Bi(III), In(III) or Ga(III) at 4 °C over a duration of 14 days. Ratio of bound metal was determined by native MS in triplicate.

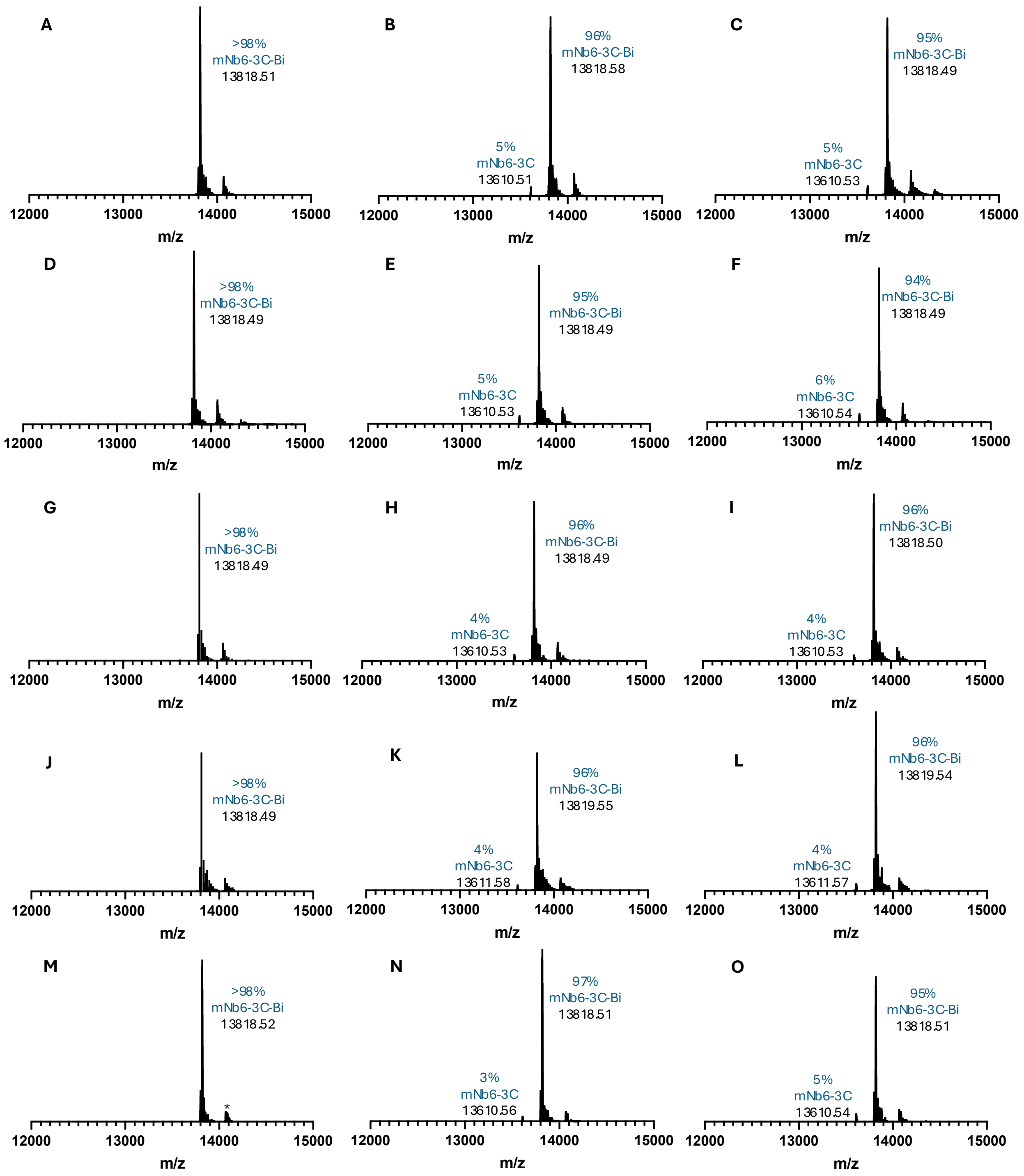


**Fig S17.** Native MS for Bi(III) bound mNb6-3C in triplicate stored at 4 °C for 0 days (A-C), 1 day (D-F), 2 days (G-I), 7 days (J-L) and 14 days (M-O).

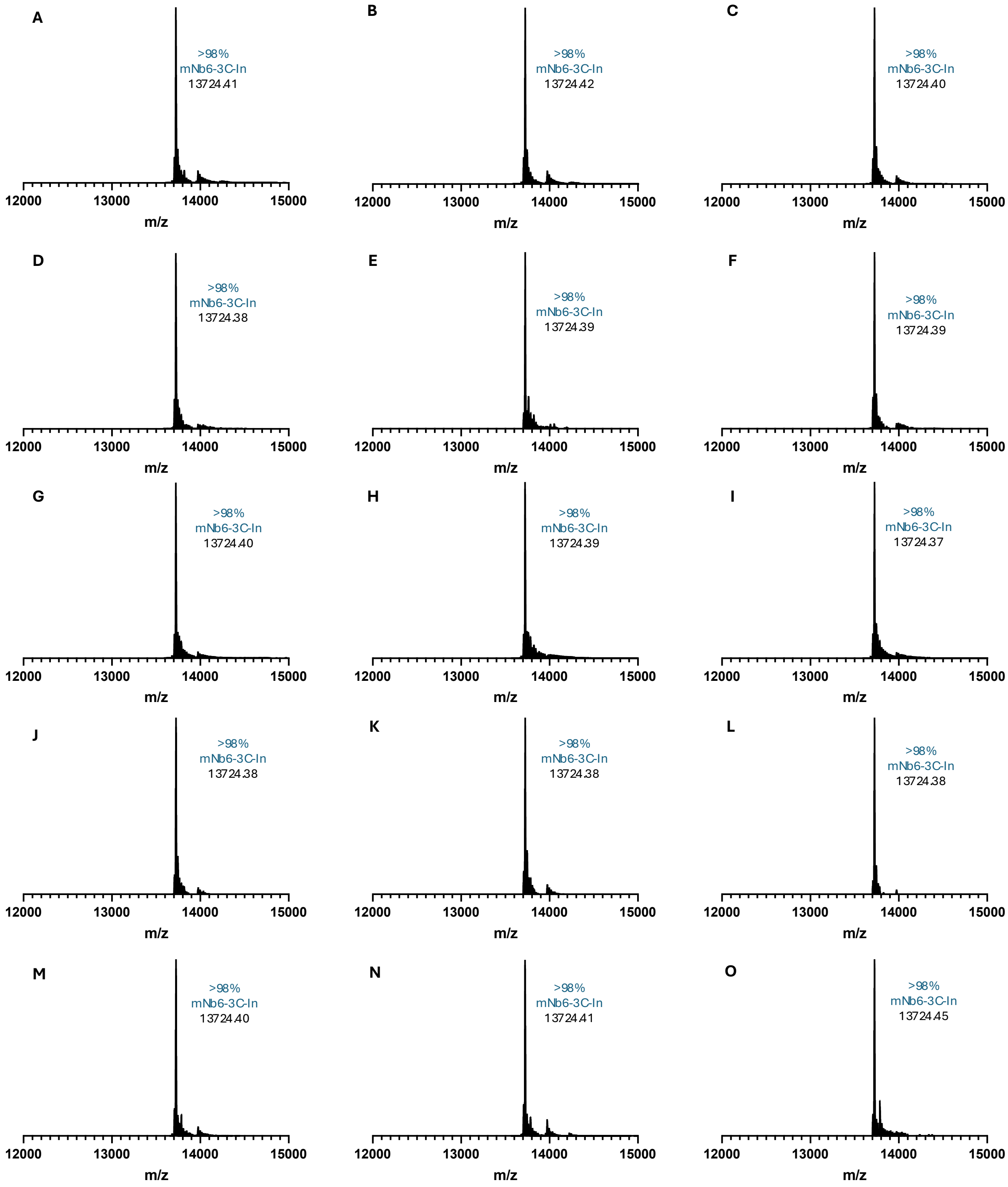


**Fig S18.** Native MS for In(III) bound mNb6-3C in triplicate stored at 4 °C for 0 days (A-C), 1 day (D-F), 2 days (G-I), 7 days (J-L) and 14 days (M-O).

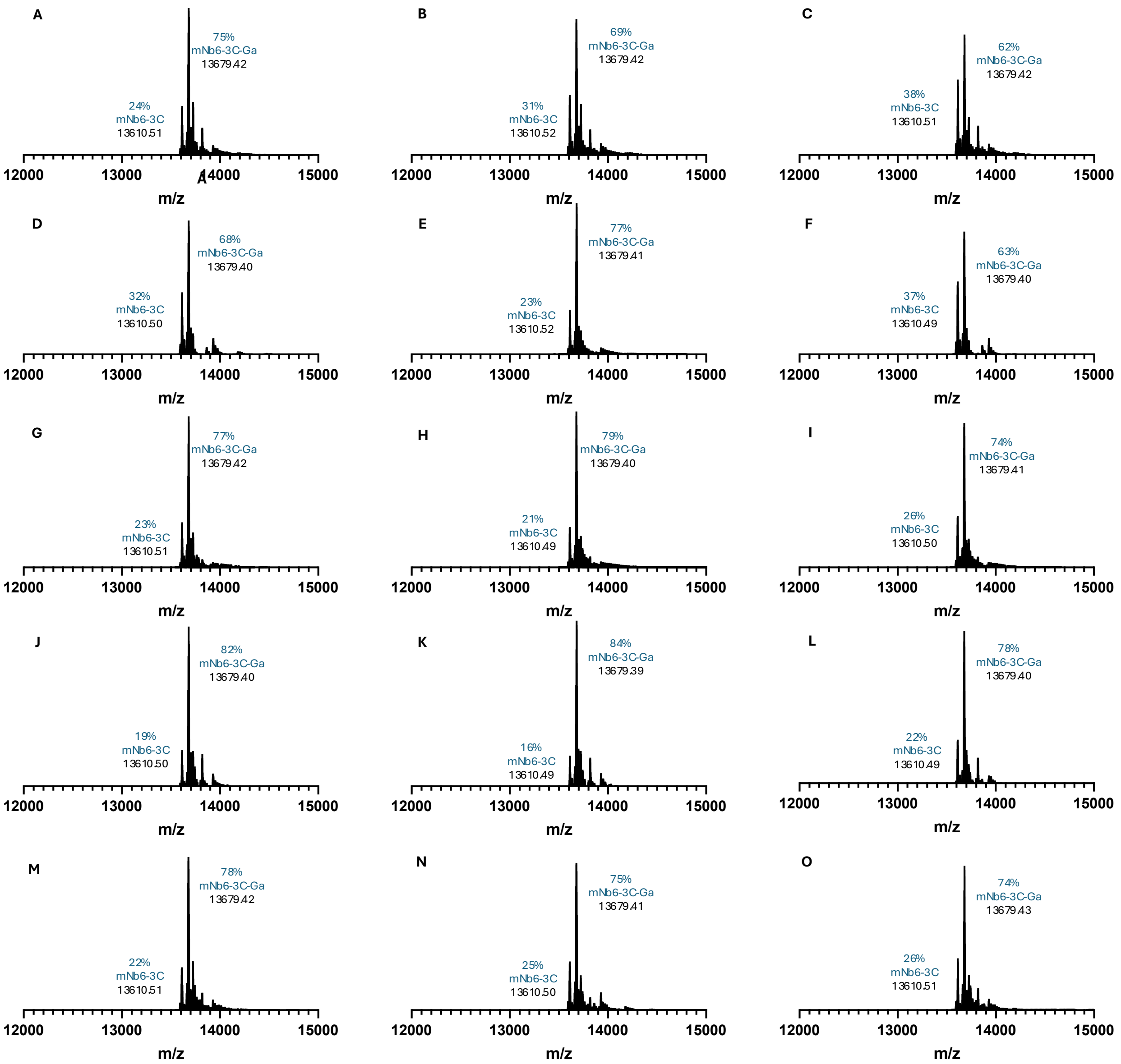


**Fig S19.** Native MS for Ga(III) bound mNb6-3C in triplicate stored at 4 °C for 0 days (A-C), 1 day (D-F), 2 days (G-I), 7 days (J-L) and 14 days (M-O).

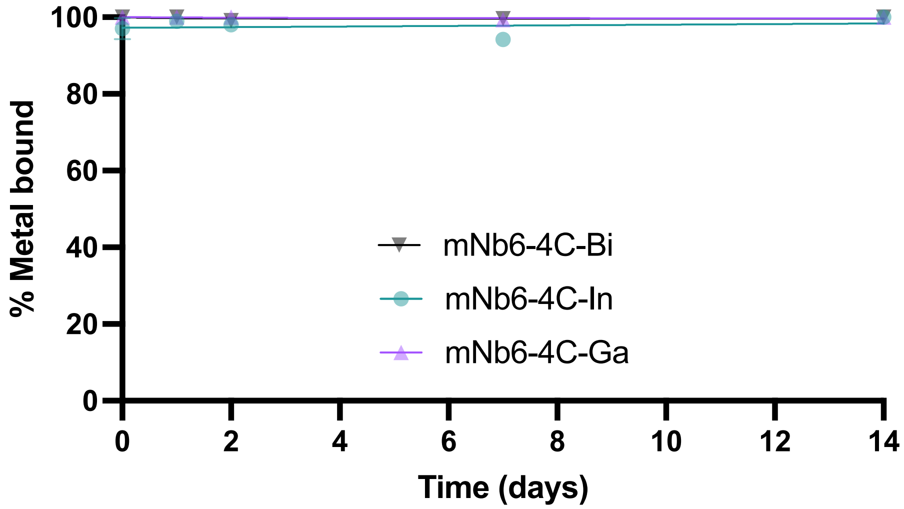


**Fig S20.** Stability of mNb6-4C in complex with Bi(III), In(III) or Ga(III) at 4 °C over a duration of 14 days. Ratio of bound metal was determined by native MS in triplicate.

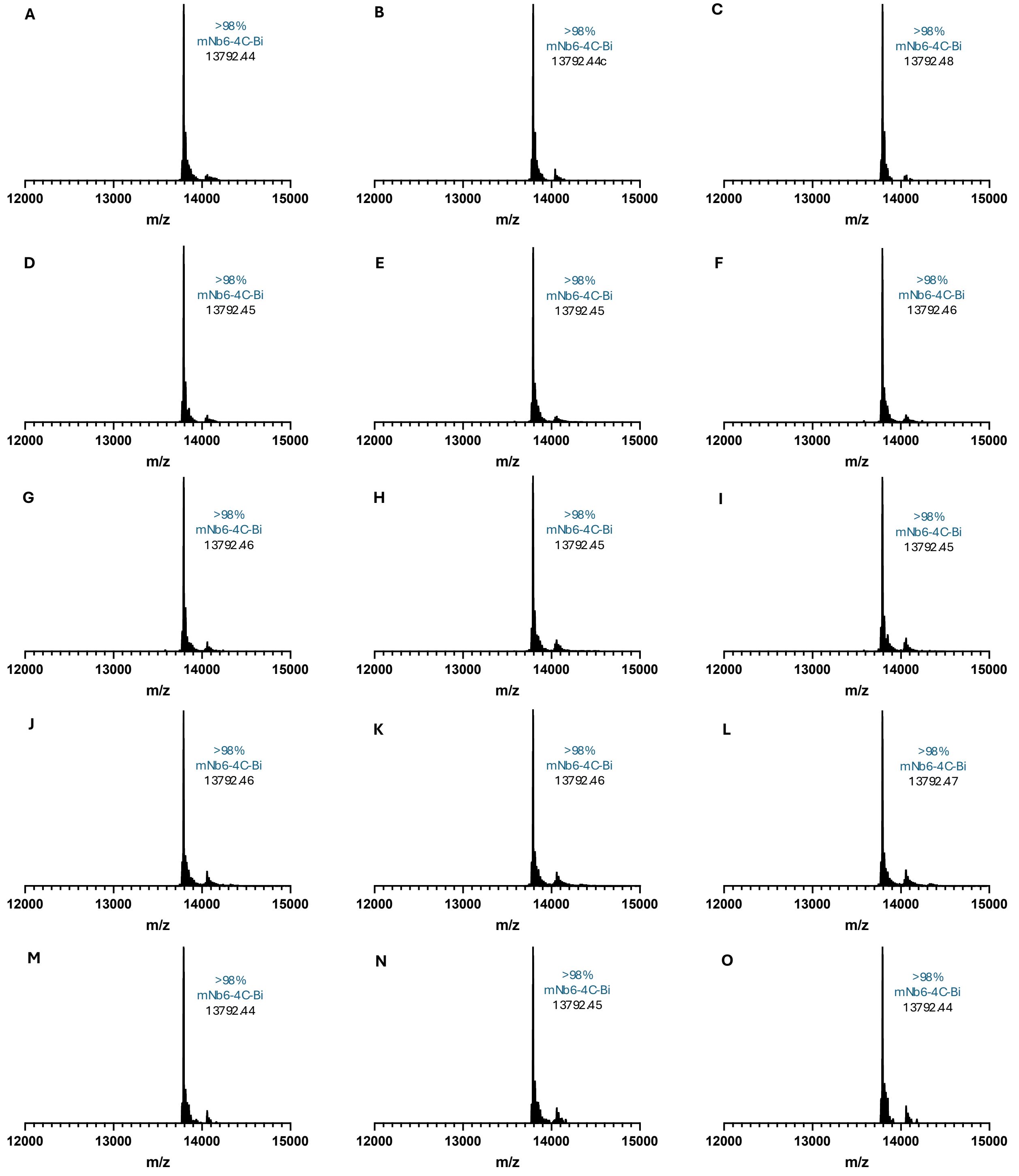


**Fig S21.** Native MS for Bi(III) bound mNb6-4C in triplicate stored at 4 °C for 0 days (A-C), 1 day (D-F), 2 days (G-I), 7 days (J-L) and 14 days (M-O).

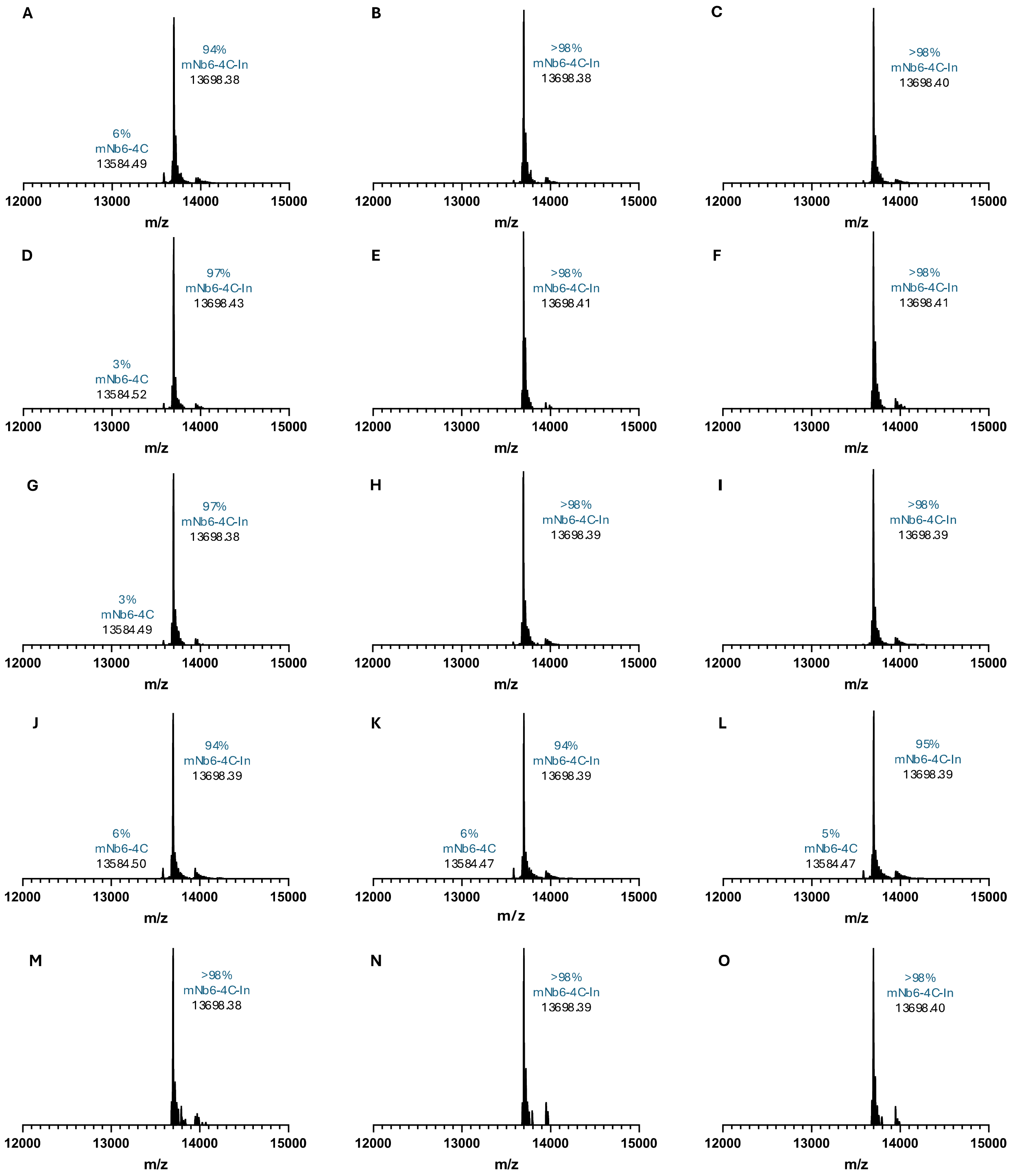


**Fig S22.** Native MS for In(III) bound mNb6-4C in triplicate stored at 4 °C for 0 days (A-C), 1 day (D-F), 2 days (G-I), 7 days (J-L) and 14 days (M-O).

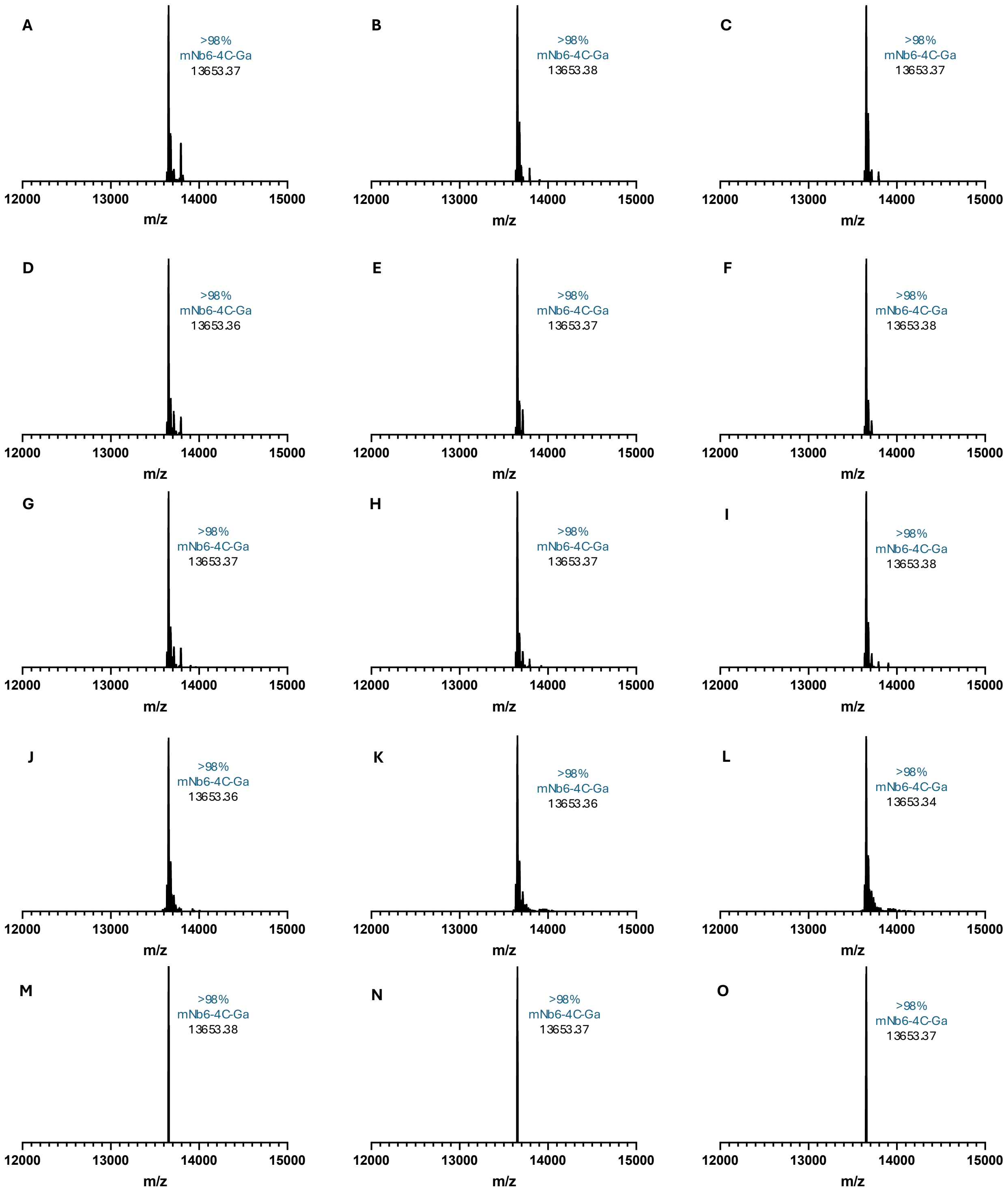


**Fig S23.** Native MS for Ga(III) bound mNb6-4C in triplicate stored at 4 °C for 0 days (A-C), 1 day (D-F), 2 days (G-I), 7 days (J-L) and 14 days (M-O).

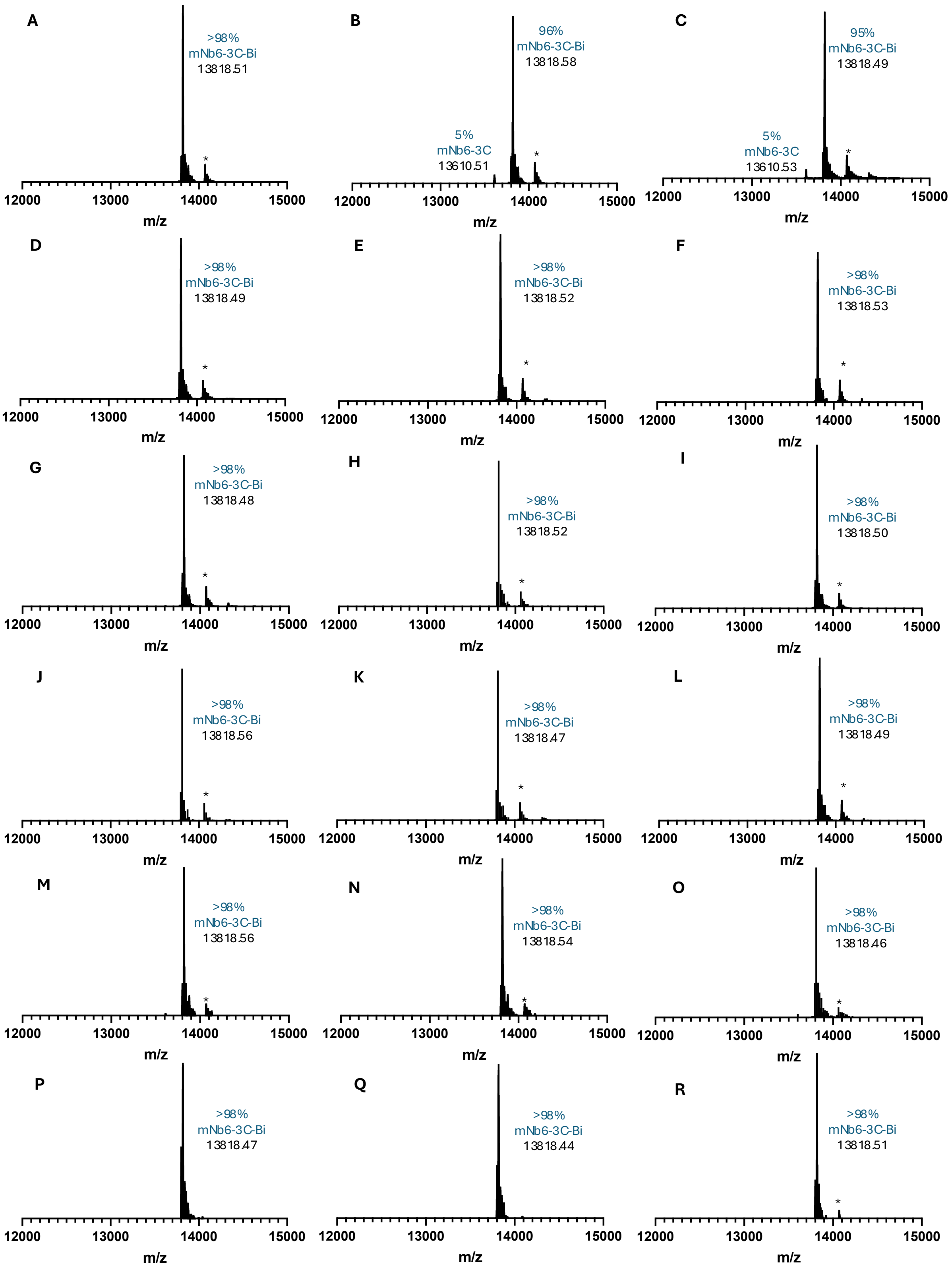


**Fig. S24.** Native MS (triplicate) of 'primed' (pre-reduced) mNb6-3C at different time points after reduction and storage at 4 °C before addition of Bi(III). Day 0 (A to C); day 1 (D to F); day 2 (G to I); day 3 (J to L); day 7 (M to O); day 14 (P to R).

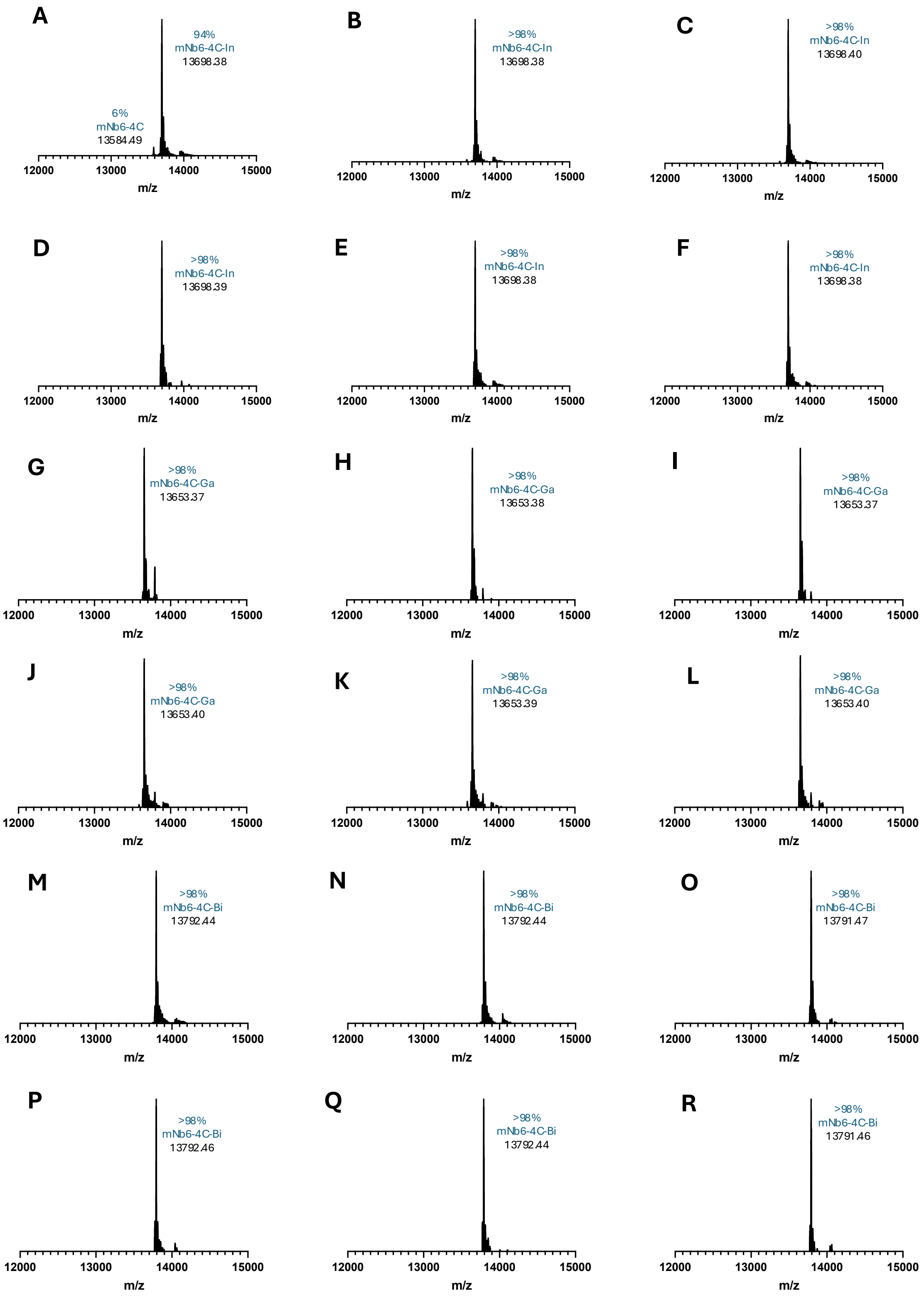


**Fig. S25.** Native MS (triplicate) of 'primed' (pre-reduced) mNb6-4C at different time points after reduction and storage at 4 °C before addition of In(III), Ga(III) or Bi(III). Day 0, In(III) (A to C); day 14, In(III) (D to F); day 0, Ga(III) (G to I); day 14, Ga(III) (J to L); day 0, Bi(III) (M to O); day 14, Bi(III) (P to R).

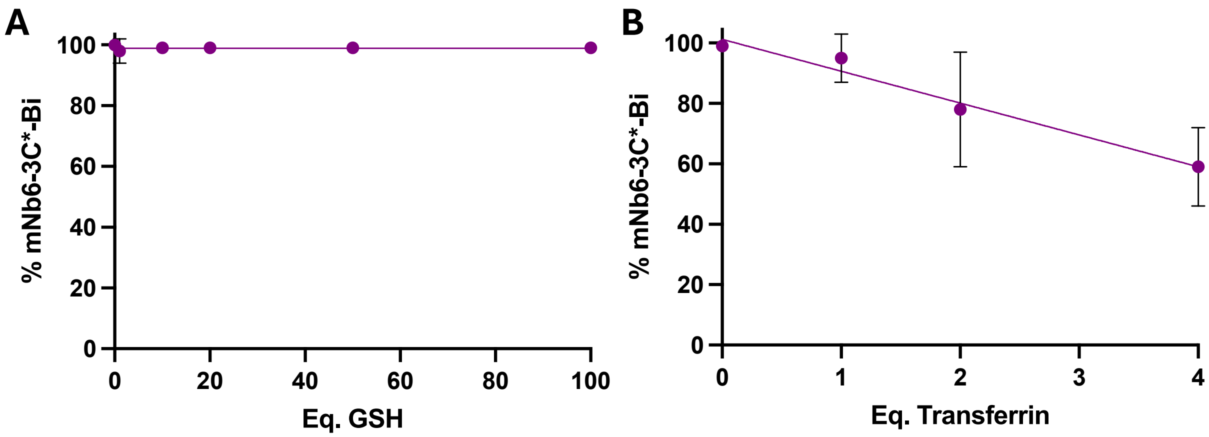


**Fig S26.** Retention of Bi(III) bound to mNb6-3C* (50 μM for C, 25 μM for D) in presence of glutathione (A) and apo-transferrin (B) after 1 h incubation at 25 °C in 100 mM ammonium acetate, pH 7.0.

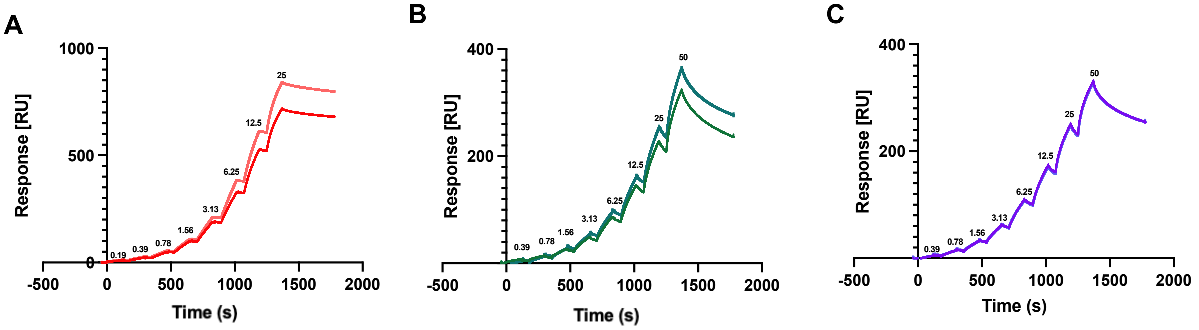


**Fig S27.** SPR sensorgrams (duplicate) showing the binding response of the SARS-CoV-2 receptor binding domain (RBD) to mNb6-3C-Bi (A), mNb6-4C-In (B), and mNb6-4C-Ga (C) immobilised on a CM5 chip in single kinetics mode. Rounded concentrations (in nM) are indicated above the response peaks.

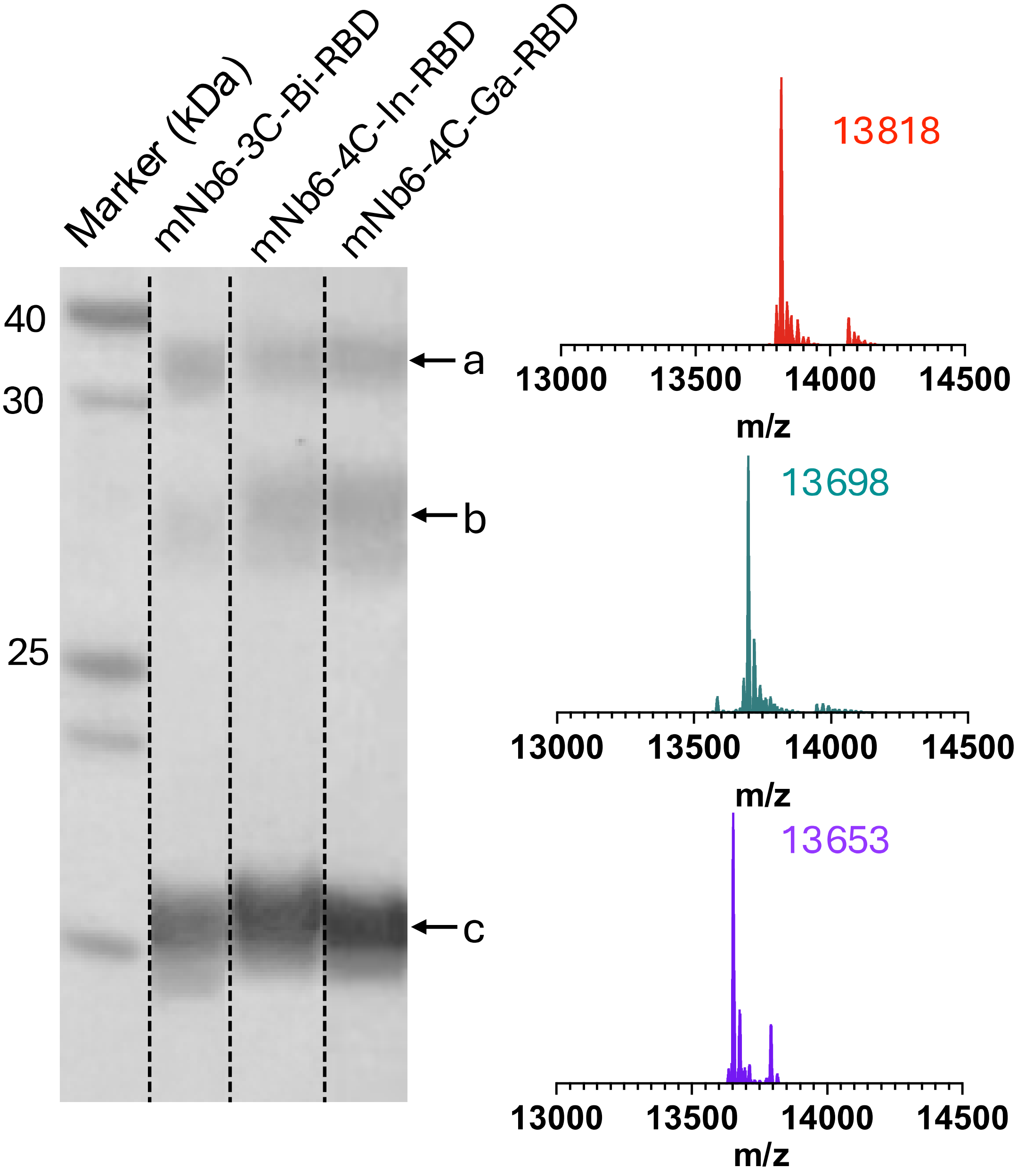


**Fig S28.** SDS PAGE of cross-linked metal-nanobody-RBD complexes. Metal-nanobodies were incubated with the SARS-CoV-2 spike RBD (1:1) for 60 min at 4 °C and subsequently crosslinked by incubation with 100 equiv. of disuccinimidyl glutarate (DSG) at 4 °C for 60 min. The samples were then loaded and run on an SDS PAGE. The native MS spectra confirm full metal coordination prior incubation (mNb6-3C-Bi, red; mNb6-4C-In, teal; mNb6-4C-Ga, purple). The mNb6:RBD complex (a), RBD (b) and metal-nanobody (c) bands are indicated.

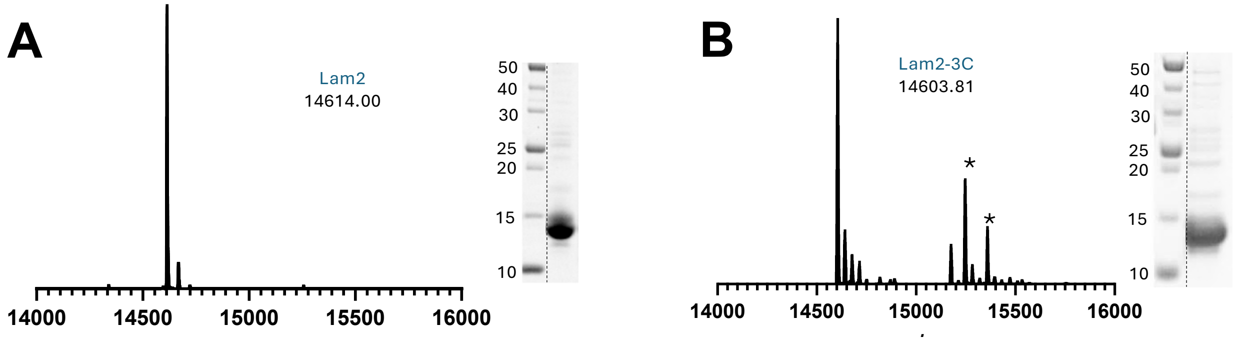


**Fig S29.** SDS-PAGE and intact protein MS confirming successful periplasmic expression of Lam2 (A) and Lam2-3C (B) in *E. coli*. Species of higher m/z ratio that correspond to a partially cleaved N-terminal leader sequence are indicated with an asterisk (*). SDS-PAGE marker proteins and their mass in kDa are shown.

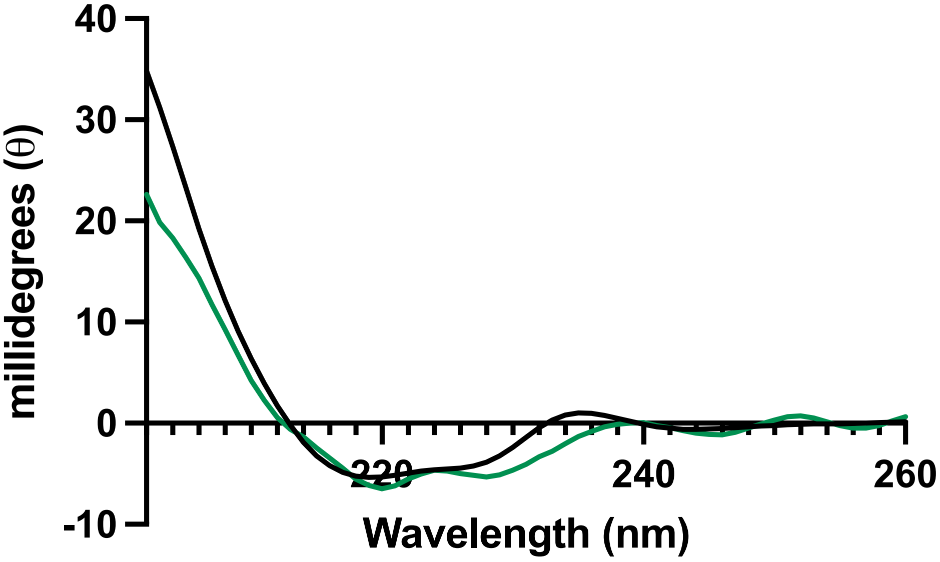


**Fig S30.** Circular dichroism (CD) spectra of Lam2 (black) and Lam2-3C-Bi (green) measured at 15 $\mu$M in 20 mM phosphate buffer, pH 7.5.

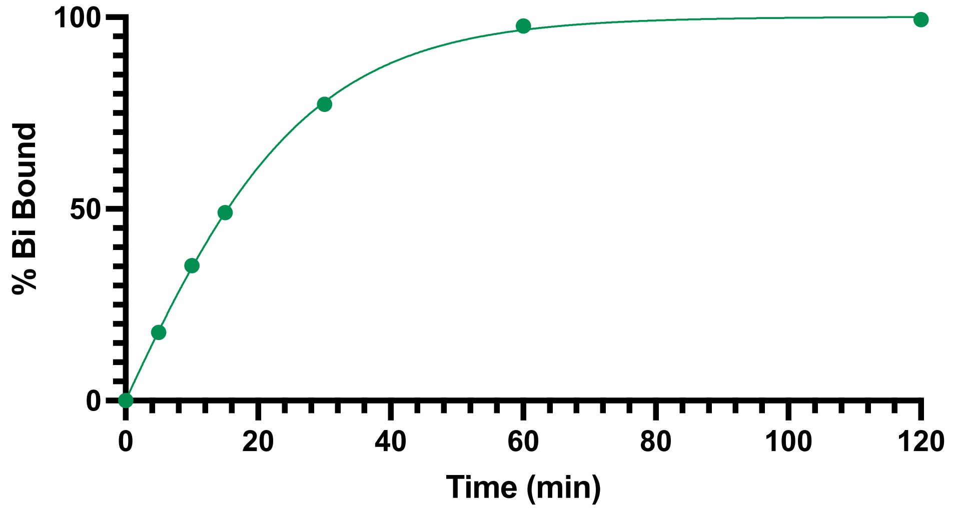


**Fig S31.** Uptake of Bi(III) by Lam2-3C in 25 mM TCEP at pH 3.0 following co-incubation with gastrodenol at 58 °C over varying durations.

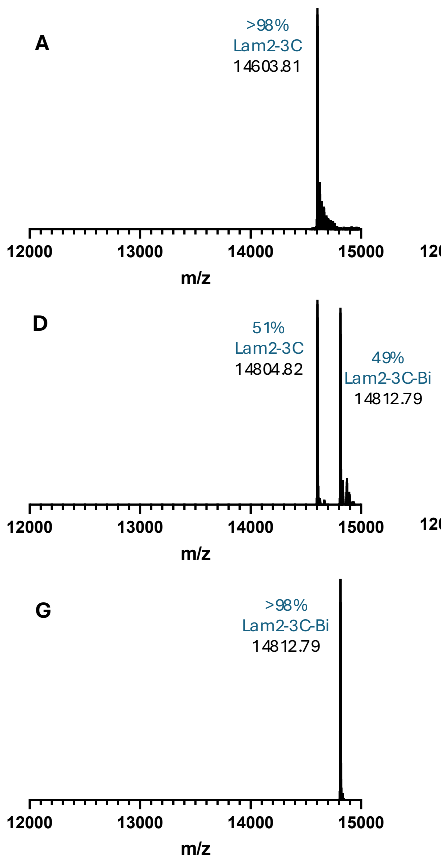


**Fig S32.** Optimization of Bi(III) uptake by Lam2-3C. Native MS of samples in 25 mM TCEP, pH 3.0 after 0 min (A), 5 min (B), 10 min (C), 15 min (D), 30 min (E), 60 min (F), and 120 min (G) co-incubation with gastrodenol at 58 °C.

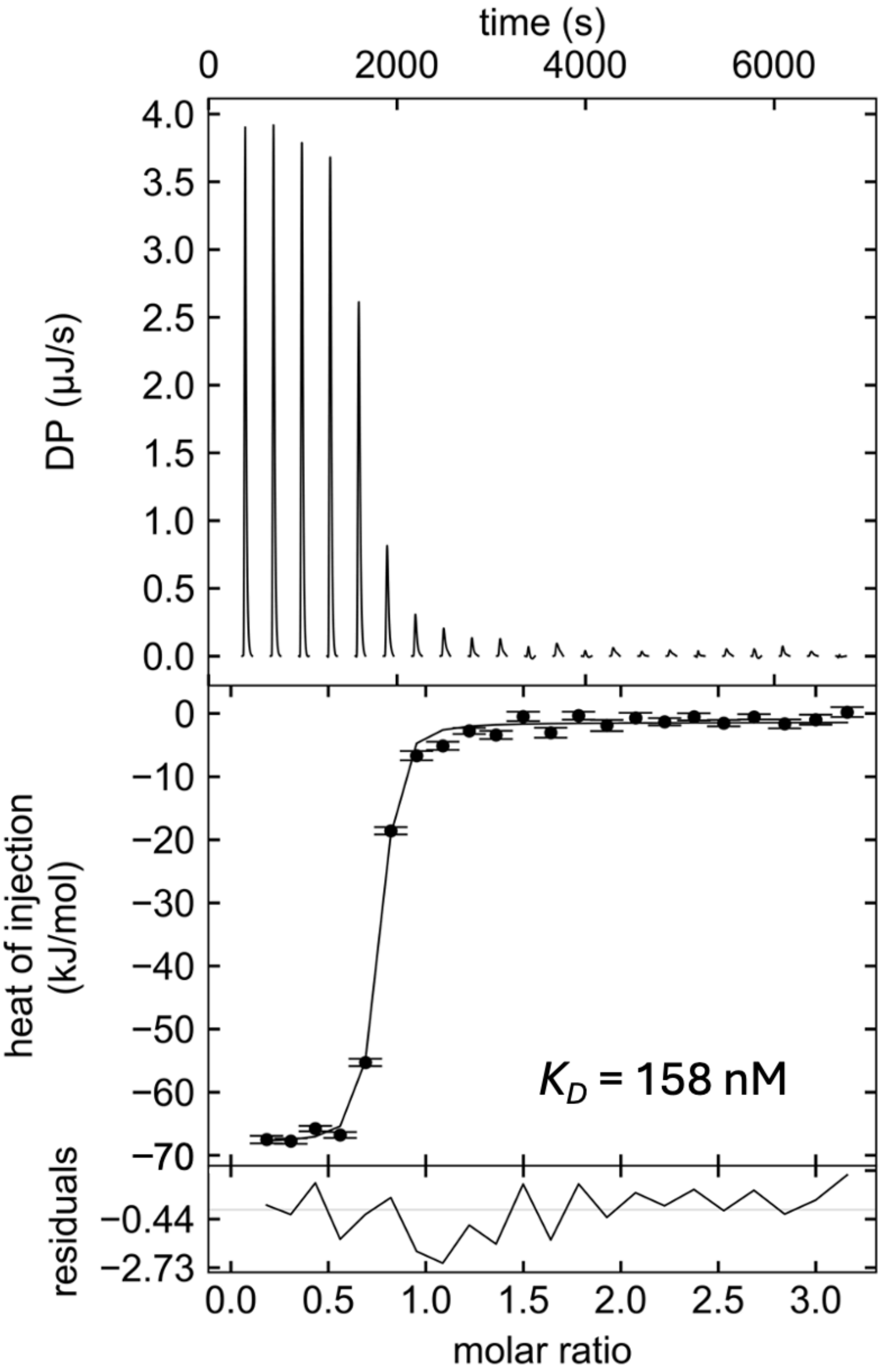


**Fig S33.** Isothermal titration calorimetry (ITC) of Lam2 and mCherry with indicated dissociation constant ($K_D$).

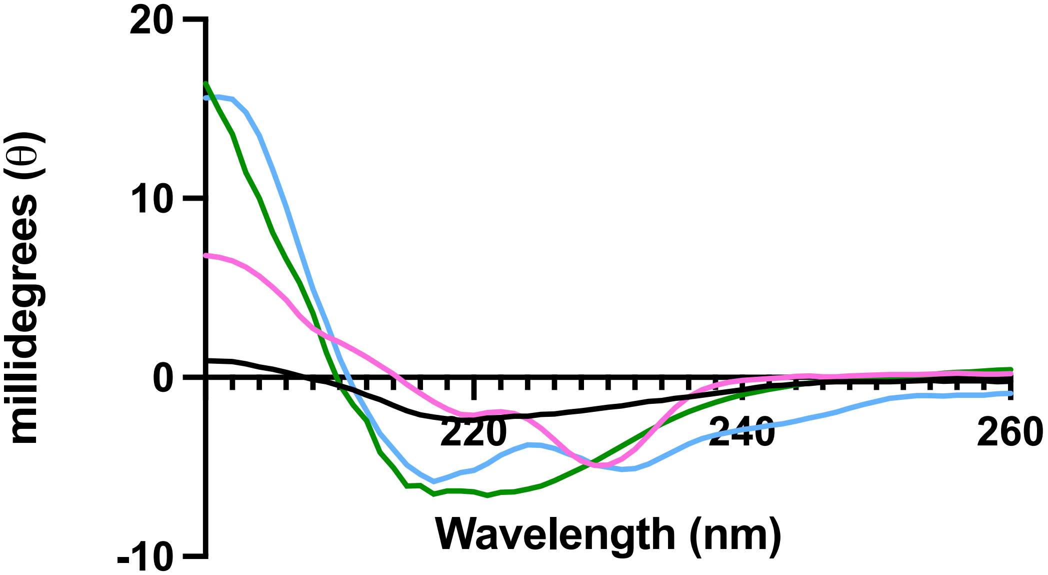


**Fig S34.** Circular dichroism (CD) spectra of 2Rs15d-5C (black), 2Rs15d-5C-Bi (pink), 2Rs15d-2A-3C (green), and 2Rs15d-2A-3C-Bi (blue) measured at 15 $\mu$M in 20 mM phosphate buffer, pH 7.5.

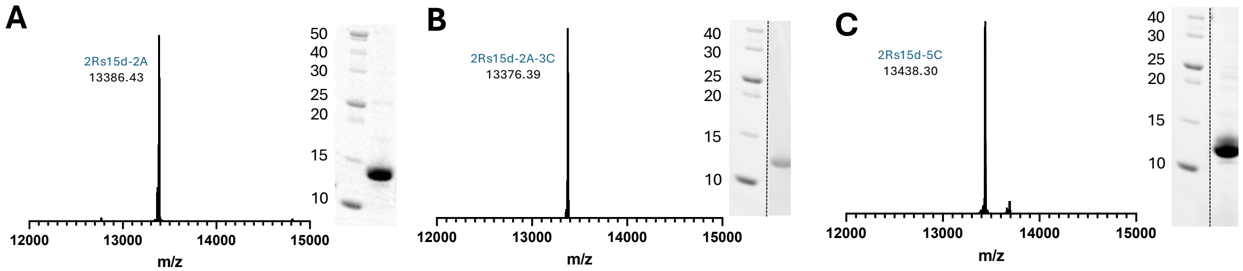


**Fig S35.** SDS-PAGE and intact protein MS confirming successful periplasmic expression of 2Rs15d-2C (A), 2Rs15d-2A-3C (B) and 2Rs15d-5C (C) in *E. coli*. SDS-PAGE marker proteins and their mass in kDa are shown.

**Table S4.** Sequences of nanobody constructs used in this study (mutations highlighted).

| Nanobody | Sequence |
|---|---|
| **mNb6** | QVQLVESGGGLVQAGGSLRLSCAASGYIFGRNAMGWYRQAPGKERELVAGITRRGSITYYADSVKGRFTISRDNAKNTVYLQMNSLKPEDTAVYYCAADPASPAYGDYWGQGTQVTVSSHHHHHH |
| **mNb6-3C** | QVQCVESGGGLVQAGGSLRLSCAASGYIFGRNAMGWYRQAPGKERELVAGITRRGSITYYADSVKGRFTISRDNAKNTVYLQMNSLKPEDTAVYYCAADPASPAYGDYWGQGTQVTVSSHHHHHH |
| **mNb6-3C*** | QVQLVCSGGGLVQAGGSLRLSCAASGYIFGRNAMGWYRQAPGKERELVAGITRRGSITYYADSVKGRFTISRDNAKNTVYLQMNSLKPEDTAVYYCAADPASPAYGDYWGQGTQVTVSSHHHHHH |
| **mNb6-4C** | QVQCVCSGGGLVQAGGSLRLSCAASGYIFGRNAMGWYRQAPGKERELVAGITRRGSITYYADSVKGRFTISRDNAKNTVYLQMNSLKPEDTAVYYCAADPASPAYGDYWGQGTQVTVSSHHHHHH |
| **Lam2** | SAQVQLVESGGGLVQAGGSLRLSCATSGFTFSDYAMGWFRQAPGKEREFVAAISWSGHVTDYADSVKGRFTISRDNVKNTVYLQMNSLKPEDTAVYSCAAAKSGTWWYQRSENDFGSWGQGTQVTVSHHHHHH |
| **Lam2-3C** | SAQVQCVESGGGLVQAGGSLRLSCATSGFTFSDYAMGWFRQAPGKEREFVAAISWSGHVTDYADSVKGRFTISRDNVKNTVYLQMNSLKPEDTAVYSCAAAKSGTWWYQRSENDFGSWGQGTQVTVSHHHHHH |
| **2Rs15d** | QVQLQESGGGSVQAGGSLKLTCAASGYIFNSCGMGWYRQSPGRERELVSRISGDGDTWHKESVKGRFTISQDNVKKTLYLQMNSLKPEDTAVYFCAVCYNLETYWGQGTQVTVSSHHHHHH |
| **2Rs15d-5C** | QVQCQESGGGSVQAGGSLKLTCAASGYIFNSCGMGWYRQSPGRERELVSRISGDGDTWHKESVKGRFTISQDNVKKTLYLQMNSLKPEDTAVYFCAVCYNLETYWGQGTQVTVSSHHHHHH |
| **2Rs15d-2A** | QVQLQESGGGSVQAGGSLKLTCAASGYIFNSAGMGWYRQSPGRERELVSRISGDGDTWHKESVKGRFTISQDNVKKTLYLQMNSLKPEDTAVYFCAVAYNLETYWGQGTQVTVSSHHHHHH |
| **2Rs15d-2A-3C** | QVQCQESGGGSVQAGGSLKLTCAASGYIFNSAGMGWYRQSPGRERELVSRISGDGDTWHKESVKGRFTISQDNVKKTLYLQMNSLKPEDTAVYFCAVAYNLETYWGQGTQVTVSSHHHHHH |

**Table S5.** Estimation of soluble mNb6-3C protein fraction after TCEP treatment at different pH.

| | | **25 mM TCEP, pH 7.5** | **25 mM TCEP pH 3.0** |
|---|---|---|---|
| Before heating at 50 °C for 15 min | $A_{280}$ | 1.35 | 1.30 |
| | Concentration (μM) | 55 | 52 |
| Post-heating at 50 °C for 15 mins | $A_{280}$ | 0.28 | 1.13 |
| | Concentration (μM) | 11.4 | 46 |

**Table S6.** Thermodynamic parameters for Lam2 and Lam2-3C-Bi binding to mCherry determined by isothermal titration calorimetry (ITC).

| | n | Δ*G* (kcal/mol) | Δ*H* (kcal/mol) | *T*Δ*S* (kcal/mol) | $K_D$ (nM) |
|---|---|---|---|---|---|
| **Lam2** | 1 | -9.3 | -15.9 | -6.6 | 158 |
| **Lam2-3C-Bi** | 1 | -10 | -12.3 | -2.3 | 46 |

**Table S7.** Expected and detected bismuth content in native PAGE samples by ICP-MS.

| Sample | Expected Bi (ppb) | Detected Bi (ppb) |
|---|---|---|
| Gel (Control) | 0 | 0.93047395 |
| Lam2-3C-Bi-mCherry | 33.4 | 24.1479996 |
| mCherry (Control) | 0 | 0.23642789 |
